# Up-regulation of cholesterol synthesis by lysosomal defects requires a functional mitochondrial respiratory chain

**DOI:** 10.1101/2024.03.06.583589

**Authors:** Francesco Agostini, Leonardo Pereyra, Justin Dale, King Faisal Yambire, Silvia Maglioni, Alfonso Schiavi, Natascia Ventura, Ira Milosevic, Nuno Raimundo

**Affiliations:** Department of Cellular and Molecular Physiology, Penn State College of Medicine, Hershey PA, USA; Department of Cellular Biochemistry, University Medical Center Goettingen, Germany; Laboratory of Systems Cancer Biology, The Rockefeller University, New York, NY, USA; IUF-Leibniz Research Institute for Environmental Medicine, Duesseldorf, Germany; Institute for Clinical Chemistry and Laboratory Diagnostic, Medical Faculty, Heinrich Heine University, Duesseldorf, Germany; Centre for Human Genetics, Nuffield Department of Medicine, University of Oxford, Oxford, UK; Penn State Cancer Institute, Penn State College of Medicine, Hershey PA, USA

## Abstract

Mitochondria and lysosomes are two organelles that carry out both signaling and metabolic roles in the cells. Recent evidence has shown that mitochondria and lysosomes are dependent on one another, as primary defects in one cause secondary defects in the other. Nevertheless, the signaling consequences of primary mitochondrial malfunction and of primary lysosomal defects are not similar, despite in both cases there are impairments of mitochondria and of lysosomes. Here, we used RNA sequencing to obtain transcriptomes from cells with primary mitochondrial or lysosomal defects, to identify what are the global cellular consequences that are associated with malfunction of mitochondria or lysosomes. We used these data to determine what are the pathways that are affected by defects in both organelles, which revealed a prominent role for the cholesterol synthesis pathway. This pathway is transcriptionally up-regulated in cellular and mouse models of lysosomal defects and is transcriptionally down-regulated in cellular and mouse models of mitochondrial defects. We identified a role for post-transcriptional regulation of the transcription factor SREBF1, a master regulator of cholesterol and lipid biosynthesis, in models of mitochondrial respiratory chain deficiency. Furthermore, the retention of Ca^2+^ in the lysosomes of cells with mitochondrial respiratory chain defects contributes to the differential regulation of the cholesterol synthesis pathway in the mitochondrial and lysosomal defects tested. Finally, we verified *in vivo*, using models of mitochondria-associated diseases in C. *elegans*, that normalization of lysosomal Ca^2+^ levels results in partial rescue of the developmental arrest induced by the respiratory chain deficiency.

## INTRODUCTION

Mitochondria and lysosomes are two fundamental organelles for metabolism. Mitochondria carry out the bulk of aerobic metabolism and have fundamental roles in Fe-S cluster synthesis, while lysosomes harbor the machinery to degrade macromolecules, obtained from the intracellular and extracellular environments, and return the building blocks to the cytoplasm and other organelles for biosynthetic reactions.

Besides their long-established metabolic functions, both of these organelles play important roles as signaling platforms, which remain an active area of investigation. Multiple signaling pathways have been implicated in the response to perturbations in mitochondrial function and dynamics, some of these acting in a tissue-specific manner (1). The mitochondrial unfolded protein response (mitoUPR) seems to integrate several aspects of the cellular response to mitochondrial stress in different tissues (2). However, some perturbations in mitochondrial functions also affect the activity of the metabolic signaling hub AMP-activated protein kinase (AMPK), depending on the specific mitochondrial defect and the tissue affected (3–7). AMPK and another signaling hub, mechanistic target of rapamycin complex 1 (mTORC1) function as respective coordinators of catabolism and anabolism (8, 9); mTORC1 promotes cellular anabolism, while AMPK represses anabolic pathways and stimulates catabolism (10, 11). mTORC1 signaling is also impacted in some instances of mitochondrial malfunction, in a defect- and tissue-specific manner (12–14). Notably, both mTORC1 and AMPK can be recruited to the lysosomal membrane for activation, thus highlighting the importance of mitochondria-lysosome coordination in the regulation of cellular metabolism (15).

Perturbations in lysosomal functions have also been shown to affect the cellular signaling hub mTORC1 (21, 22). Further, acidification of the lysosomal lumen is needed for the activity of the hydrolases and for cholesterol efflux from lysosomes to endoplasmic reticulum (ER) (16, 17). Loss of lysosomal acidification results in functional iron deficiency and pseudo-hypoxia signaling (18–20).

Importantly, mitochondria and lysosomes are interdependent (23, 24). Impairment of mitochondrial respiratory chain and/or mitochondrial dynamics results in dysfunctional lysosomes, with excessive storage of undigested material in the lysosomal lumen (7, 25, 26). Lysosomal defects in sphingolipid catabolism, cholesterol trafficking and impaired lysosomal acidification result in repression of the transcriptional program of mitochondrial biogenesis and impaired respiratory chain function (27). The mitoUPR signaling in response to mitochondrial malfunction is partially mediated by lysosomes (28, 29). Furthermore, mitochondria and lysosomes physically interact with each other via membrane contact sites, as well as with other organelles, especially the ER (30, 31). The contact sites between mitochondria and lysosomes have been linked to the ability of lysosomes to regulate mitochondrial dynamics by stimulating mitochondrial fission (32).

While there is negative feedback between mitochondrial malfunction and lysosomal malfunction, it is still not clear how defective mitochondria and lysosomes impact other cellular processes, in particular lipid homeostasis. Here, we use bulk cellular transcriptome data to unveil the signaling pathways that are affected by mitochondrial and lysosomal malfunction, and show that these two organelles have opposite impacts on the synthesis of lipids, particularly on SREBF1-dependent cholesterol synthesis, and that these effects are independent of AMPK and mTORC1 signaling. These results are relevant to understand the complexity of organelle crosstalk and its interplay with cellular metabolism.

## RESULTS

### Mitochondria and lysosomal defects have opposite effects on regulation of cholesterol synthesis

Mitochondria and lysosomes show functional interdependence: the malfunction of one leads to a malfunction in the other. Because both organelles have signaling roles, we sought to determine which aspects of cellular signaling and metabolism are affected both by mitochondria and lysosomal dysfunction. To study the signaling environment in a broad and unbiased manner, we characterized the cellular transcriptome using bulk RNA sequencing. As models of organelle dysfunction, we employed several human and murine cellular models of chronic mitochondrial and lysosomal malfunction (see **Figure 1A**). As a model of mitochondrial malfunction we used complex III deficiency, caused by the stable knock-down of complex III subunit UQCRC1, through lentivirus-delivered shRNA in HeLa cells, henceforth referred as UQCRC1-kd (7). We previously showed that these cells are an adequate model of mitochondrial respiratory chain deficiency, with decreased O_2_ consumption and increased superoxide levels (7). As models of lysosomal malfunction, we used HeLa cells with a stable silencing of acid alpha-glucosidase (GAA-kd) or of cathepsin B (CTSB-kd), also obtained by shRNA-mediated silencing (**Supplementary Figure 1**). GAA is the enzyme responsible for the hydrolysis of glycogen to glucose in the lysosomal lumen, and cathepsin B is a member of the calpain-like family of cysteine lysosomal proteases. Both GAA and CTSB are located to the lysosomal lumen.

**Figure 1.**
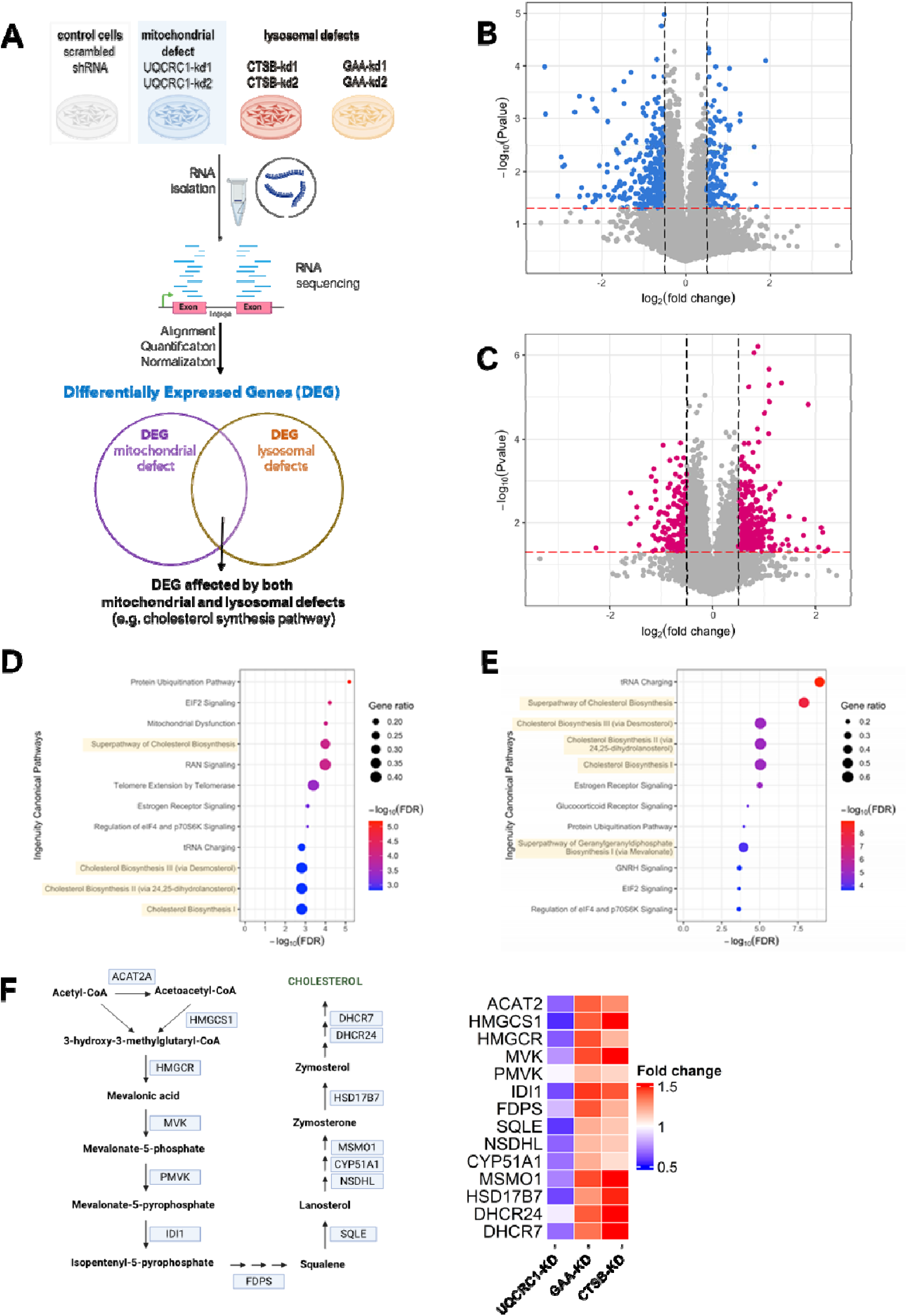
Defects in mitochondria and lysosomes have opposite effects on the regulation of cholesterol synthesis pathway. **(A)** Schematic representation of the RNAseq analysis workflow, in which one mitochondrial (UQCRC1-kd) and two lysosomal (CTSB-kd and GAA-kd) dysfunction models were examined. **(B,C)** Volcano plots of the transcriptome data analyses showing the differentially expressed genes between UQCRC1-kd and control HeLa lines (B), and those differentially expressed between the CTSB-kd and GAA-kd lines and the controls (C). **(D)** Canonical pathways altered in the mitochondrial dysfunction cell model and **(E)** in the lysosomal defective cell models. The cholesterol biosynthesis super-pathways are shadowed. Associated P values were determined according to analysis in the database for annotation, visualization, and integrated discovery (Fisher exact P-value). **(F)** Representation of the cholesterol synthesis pathway indicating the genes coding the enzymes, and heatmaps showing the respective transcripts levels (fold change) in UQCRC1-kd, GAA-kd and CTSB-kd cells.

For each gene silenced, we used two clones of HeLa cells, each made with a different shRNA, and two clones with scrambled shRNA as controls. First, we identified the differentially expressed transcripts between two UQCRC1-kd lines and the two control lines, and those differentially expressed between the lysosomal lines and the controls. For the mitochondrial defects we obtained 2116 differentially expressed genes (DEG) between UQCRC1-kd and scrambled control, of which 501 were up-regulated in UQCRC1-kd and 1615 down-regulated (**Figure 1B**). The lysosomal defects models yielded 2877 DEG, of which 994 were up-regulated in the lysosomal defect group (comparing all lines of GAA-kd and CTSB-kd versus the two controls) and 1883 down-regulated (**Figure 1C**). The DEG lists are presented in Supplementary Table I.

Curiously, when we subjected these DEG lists to pathway analysis using the Ingenuity Pathway analysis software (Qiagen), the most enriched pathways in both the mitochondrial and lysosomal defects were all related with the synthesis of mevalonate and of cholesterol (**Figure 1D-1E**). Yet, while the transcript levels of cholesterol synthesis enzymes were globally up-regulated in the lysosomal defects, they were globally down-regulated in the mitochondrial defects (**Figure 1F**).

To confirm this interesting observation, we isolated RNA from a different batch of UQCRC1-kd and control cells, and measured the transcript levels of cholesterol synthesis enzymes by qPCR. In agreement with the RNAseq results, we observed that the transcript levels of cholesterol synthesis enzymes are globally down-regulated in UQCRC1-kd cells (**Figure 2A**). To test if different perturbations of the respiratory chain would have the same impact on cholesterol synthesis, we used embryonic fibroblasts isolated from mice lacking a subunit of complex I, NDUFS4 (NDUFS4-KO) and the corresponding wild-type (WT) littermate controls. In these murine cells, similarly to the human cells, the transcript levels of cholesterol synthesis enzymes were also down-regulated (**Figure 2B**), supporting a transcriptional down-regulation of this pathway in response to impaired respiratory chain.

**Figure 2.**
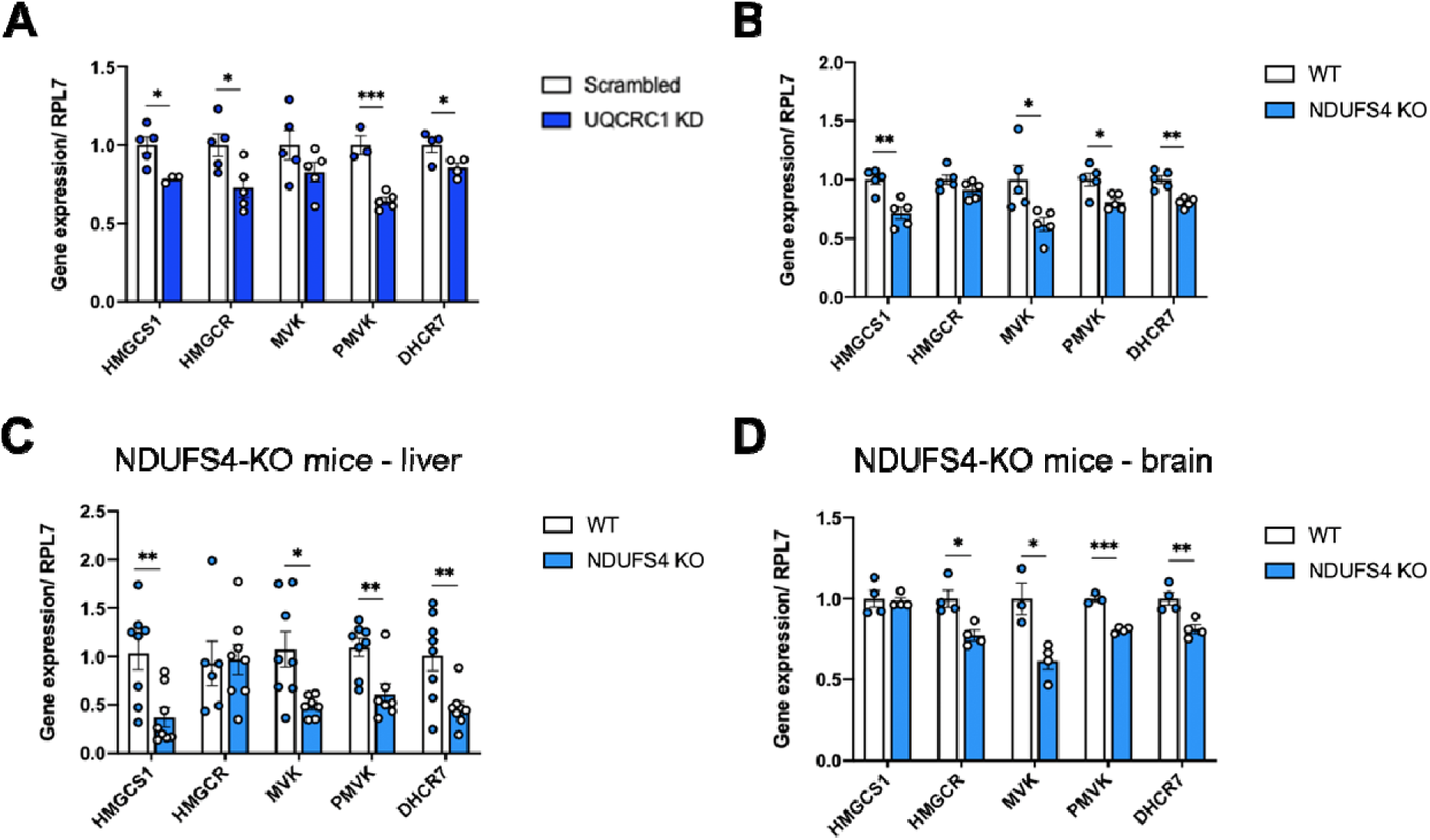
Expression of cholesterol synthesis enzymes is inhibited in cells and organs with mitochondrial respiratory chain deficiency. Real-time qPCR of genes involved in the cholesterol synthesis pathway performed in control and UQCRC1-kd HeLa cells **(A)**, NDUFS4 WT and KO MEF cells **(B)**, and in the livers and brains of NDUFS4 WT and KO mice **(C,D)**. At least 4 independent samples/animals were evaluated. Statistical significance between control and mutant samples for each gene was determined by unpaired t-test, * p < 0.05, ** p < 0.01, *** p < 0.001.

Next, we examine if our findings in cultured cells were also valid *in vivo*. We used the liver of NDUFS4-KO mice and their WT littermates and observed that the transcript levels of cholesterol synthesis enzymes were globally decreased in NDUFS4-KO liver (**Figure 2C**). Similar results were obtained in brains isolated from NDUFS4-KO mice (**Figure 2D**). Altogether, these results show that chronic perturbations in mitochondrial respiratory chain lead to a down-regulation of the transcript levels of cholesterol synthesis enzymes, both in cultured cells and in multiple mouse organs.

Next, we sought to validate the transcriptome results obtained with the cells harboring lysosomal defects. We tested new vials of HeLa GAA-kd and CTSB-kd, and again observed an up-regulation of the transcript levels of cholesterol synthesis enzymes by qPCR (**Figure 3A**). We also examined mouse fibroblasts lacking lysosomal proteins, namely GAA-KO (**Figure 3B**), CTSL-KO (**Figure 3C**), CTSB-KO (**Figure 3D**) and LAMP2-KO (**Figure 3E**), and in all cases we observed a robust up-regulation of the transcript levels of cholesterol synthesis enzymes, although to varying degrees of magnitude. To test if these results could also be observed *in vivo*, we measured the transcript levels of cholesterol synthesis enzymes in the liver of GAA-KO mice, and observed an increase in the expression of at least two enzymes (**Figure 3F**). These results show that impairment of lysosomal function results in a transcriptional up-regulation of cholesterol synthesis enzymes both in cultured cells and *in vivo* organs. Of note, these results are consistent with the transcriptional up-regulation of cholesterol synthesis upon treatment with an inhibitor of the lysosomal v-ATPase, bafilomycin A1 (18, 33, 34).

**Figure 3.**
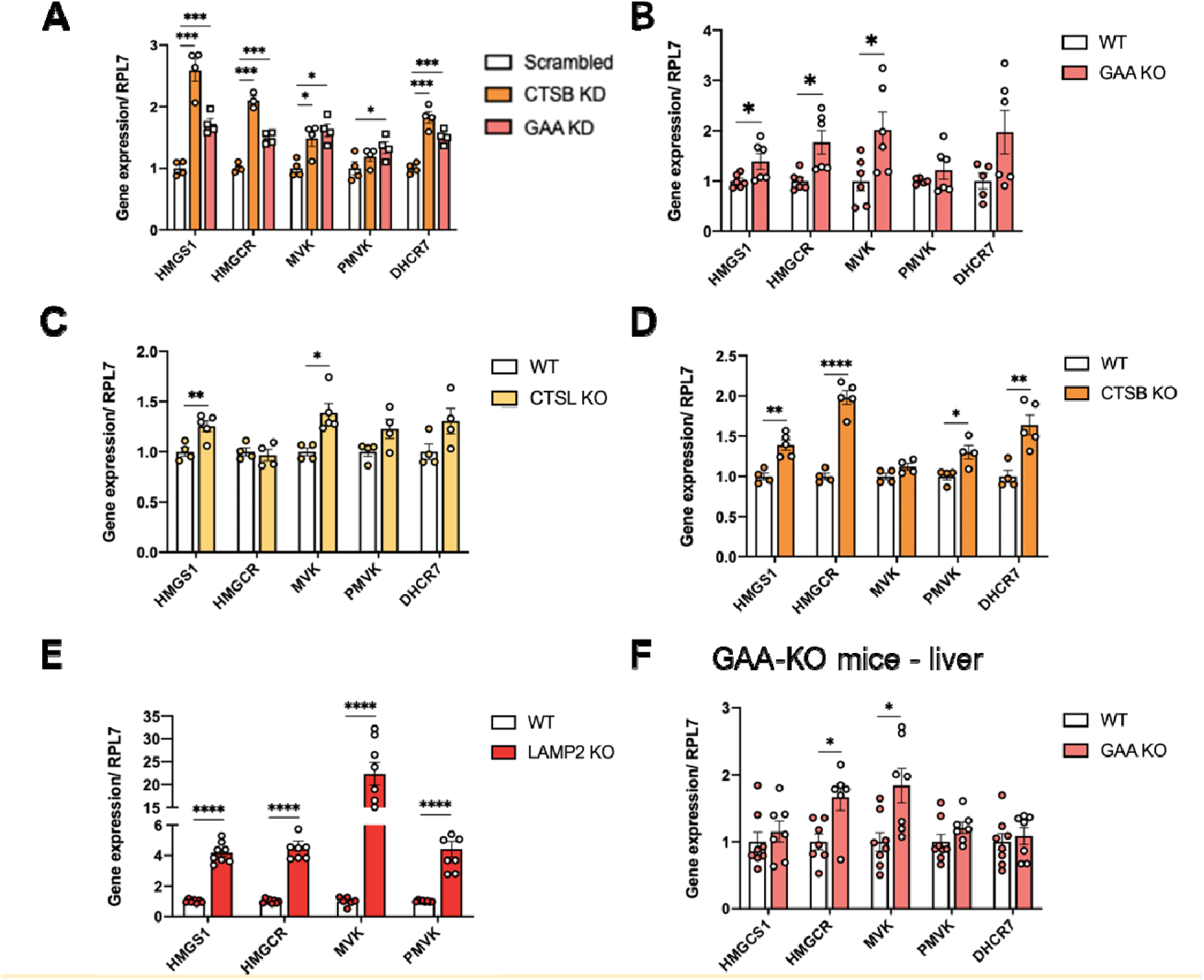
Transcriptional control of cholesterol synthesis is upregulated in cells and organs with lysosomal defects. **(A)** Real-time qPCR showing the transcript level of cholesterol synthesis enzymes in CTSB-kd, GAA-kd and control HeLa cells. At least 3 independent samples were evaluated. Statistical significance between control and mutant samples for each gene was determined by two-way ANOVA with Tukey’s multiple comparisons test. * p < 0.05, ** p < 0.01, *** p < 0.001. Th same set of transcripts was analyzed via RT qPCR in knockout MEFs for GAA **(B)**, CTSL **(C)**, CTSB **(D)**, LAMP2 **(E)** and relative WT controls, as well as in the livers of GAA WT and KO mice **(F)**. At least 4 independent animals were evaluated. Unpaired t-test was used to determine the statistical significance between control and KO samples for each gene. * p < 0.05, ** p < 0.01, *** p < 0.001, **** p < 0.0001.

It is known that lysosomal cholesterol egress requires the joint action of the proteins NPC2 and NPC1, as well as acidic pH in the lysosomal lumen (34). While a population of lysosomes is unable to acidify properly in cells lacking GAA (18, 35), there is no indication that loss of CTSB or CTSL impair lysosomal acidification, although these proteases may be involved in NPC1 maturation (36). Furthermore, lysosomal pH is increased (neutral) in the UQCRC1-kd cells, which also present lysosomal Ca^2+^ accumulation and inhibited lysosomal hydrolysis (7), suggesting that lysosomal impairment only leads to increased expression of cholesterol synthesis enzymes in the presence of a functional mitochondrial respiratory chain. We therefore sought to further explore the mechanisms underlying the differential regulation of the expression of cholesterol synthesis enzymes under mitochondrial and lysosomal defects.

### Downregulation of cholesterol synthesis pathway in cells with mitochondrial malfunction is associated with decreased SREBF1 processing and activity

The synthesis of cholesterol is regulated at transcript level by two transcription factors, SREBF1 and SREBF2 (37). These proteins are tethered to the ER membrane through interactions with SCAP (37), in a cholesterol-dependent manner: high levels of cholesterol in the ER membrane retain the SCAP/SREBF1 or SCAP/SREBF2 complexes at the ER, while a decrease in ER cholesterol content releases them towards further activation (37). The retention of SCAP/SREBF1 or SCAP/SREBF2 complexes at the ER membrane is also stimulated by two proteins, INSIG1 and INSIG2. After their release from the ER, they are further processed at the Golgi by two proteases S1P and S2P, and released in their mature form to translocate to the nucleus, where they bind the promoters of their target genes and stimulate transcription (37). Given that the effect of UQCRC1-kd and NDUFS4-KO on the transcriptional regulation of cholesterol synthesis is similar, we used the NDUFS4-KO cells to follow-up mechanistically. We observed that in NDUFS4-KO whole cell extracts, mature SREBF1 was decreased (**Figure 4A**), while SREBF2 protein levels were similar in NDUFS4-KO and WT cells (**Figure 4B**). The transcript levels of SREBF1 and SREBF2 were not changed in the NDUFS4-KO cells (**Supplementary Figure 2A**), suggesting that the reason underlying the decrease in SREBF1 protein levels is post-transcriptional. To further explore why SREBF1 levels are decreased in NDUFS4-KO cells, we briefly treated the cells with the proteasome inhibitor MG132 (2h, 10µM), to determine if ubiquitin-mediated proteasomal degradation may be responsible. SREBF1 can be degraded by the proteasome when it is phosphorylated at Ser73 by the kinase GSK-3B, which allows the ubiquitin ligase FBXW7 to recognize SREBF1 and ubiquitinate it (38). Curiously, in MG132-treated NDUFS4-KO cells, the levels of SREBF1 were returned to the levels present in WT cells (**Figure 4A**), while no changes were observed in SREBF2 levels (**Figure 4B**). These results suggest that in cells with respiratory chain deficiency, SREBF1 is continuously degraded, in a proteasome-dependent manner.

**Figure 4.**
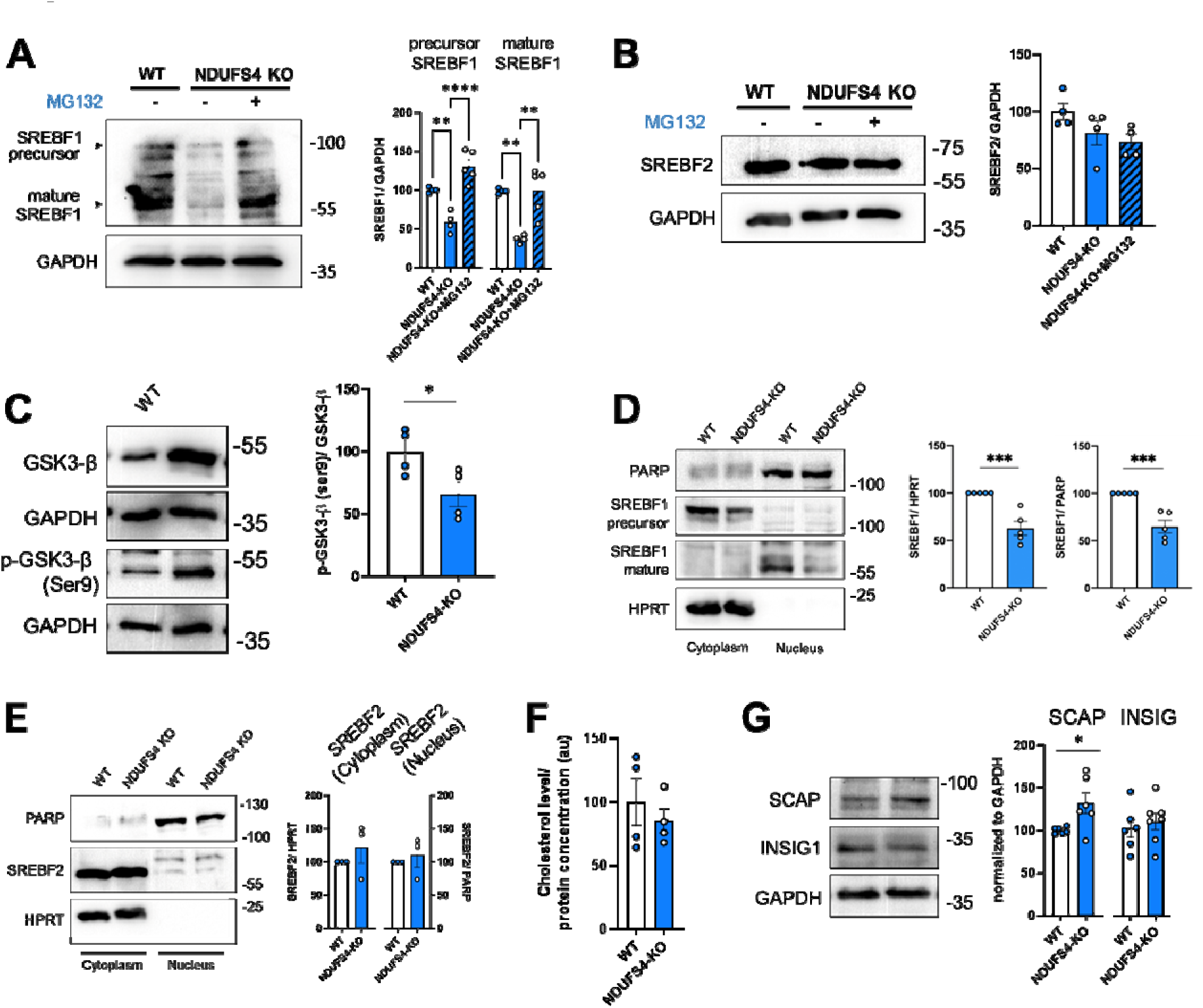
Mitochondrial defects impair the cholesterol pathway by altering the SREBF1 processing and activity. **(A,B)** Western blot analyses of SREBF1 (A) and SREBF2 (B) in WT and NDUFS4-KO fibroblasts, untreated and treated with the proteasome inhibitor MG132 (2 h, 10µM). GAPDH was used as the loading control. At least 4 independent experiments were evaluated. Statistics: one-way ANOVA with Tukey’s multiple comparisons test, * p < 0.05, ** p < 0.01, **** p < 0.0001. **(C)** Western blot analysis showing the total form of GSK3B total and its phosphorylation at the Ser9. GAPDH was used as loading control. 4 independent samples were evaluated and unpaired t-test was used for statistical analysis. *** p < 0.001 **(D,E)** Western blots of nuclear vs. cytosolic fractions from WT and NDUFS4-KO fibroblasts probed with anti-SREBF1 (D), anti-SREBF2 (E), anti-PARP (nuclear marker) and anti HPRT (cytosolic marker) antibodies. At least 4 independent experiments were done. Statistics: unpaired t-test, *** p < 0.001. **(F)** Cholesterol level measured in the ER fractions isolated from WT and NDUFS4-KO MEFs. The level of cholesterol was normalized by the total amount of protein in the corresponding fraction. 4 independent samples were analyzed. Statistical analysis was performed using unpaired t-test. **(G)** Western blot analysis of SCAP and INSIG1 levels in WT and NDUFS4-KO fibroblasts. At least 6 independent samples were evaluated. Unpaired t-test, * p < 0.05.

This result also implies that GSK3B activity may be increased in response to respiratory chain impairment. Therefore, we probed the levels of GSK3B in NDUFS4-KO whole cell extracts (**Figure 4C**) as well as the phosphorylation status of Ser9/21 (both inhibitory of GSK3B when phosphorylated). The ratio between GSK3B-Ser9/21-P and total GSK3B was decreased (**Figure 4C**), which suggests that the activity of GSK-3B is increased in cells with impaired mitochondrial respiratory chain.

While the levels of SREBF1/2 immature precursors may impact their activity, the key regulatory step is maturation. Because the activity of mature SREBF1/2 is associated with its nuclear localization, we next prepared nuclear extracts from WT and NDUFS4-KO cells and observed a decrease in SREBF1 levels in the nucleus of NDUFS4-KO fibroblasts (**Figure 4D**), supporting the decreased activity of SREBP1 in these cells. We also observed that nuclear levels of SREBF2 in NDUFS4-KO cells were similar to WT (**Figure 4E**). Further, the levels of mature SREBF1 were increased in cellular models of lysosomal defects, namely mouse fibroblasts lacking CTSB and CTSL (**Supplementary Figure 2B**).

As mentioned above, the levels of cholesterol in the ER membranes determine if SREBF1/2 remain sequestered by SCAP at the ER membrane or are released and further processed to promote its nuclear localization and transcriptional activity. This could be explained by perturbation of the levels of cholesterol in the ER in cells with mitochondrial defects (higher ER cholesterol levels) and lysosomal defects (lower ER cholesterol levels). We therefore tested if the levels of ER cholesterol were altered in cells with respiratory chain deficiency. Using a biochemical procedure, we isolated ER (**Supplementary Figure 2C-D**) and using the ER-rich fractions we measured the levels of cholesterol. We observed that there was no difference in the amount of cholesterol in ER membranes from WT and NDUFS4-KO cells (**Figure 4F**). Given that there are no changes in cholesterol concentration in the ER that can explain the lower levels of mature SREBF1 in NDUFS4-KO cells, we tested if the proteins that process the maturation of SREBF1 might be affected in the NDUFS4-KO cells. We observed that the protein levels of SCAP are increased in NDUFS4-KO cells relative to WT, while the levels of INSIG are not changed (**Figure 4G**), implying that the decreased maturation of SREBF1 is not due to lower levels of its processing proteins.

Altogether, these results show that the levels of SREBF1 precursor protein, and consequently activity of mature SREBF1 are decreased in the models of respiratory chain deficiency, while increased SREBF1 processing and activity occur in the lysosomal malfunction models. Given that the cells with respiratory chain deficiency also have defective lysosomes, the inhibition of the respiratory chain seems to modify the effect of lysosomal malfunction on cholesterol homeostasis, involving the GSK-3B-mediated degradation of SREBF1 in cells with mitochondrial respiratory chain impairments.

### The mechanisms of cholesterol trafficking and homeostasis are fully functional in cells with mitochondrial malfunction

To test if other mechanisms besides SREBF1 regulation were affecting cholesterol homeostasis in cells with respiratory chain deficiency, we probed cholesterol trafficking in these cells. Cholesterol can reach the ER via lysosomes or via plasma membrane (39). Egress of cholesterol from the lysosomal lumen and its trafficking to the ER is a key process affected by loss of lysosomal acidification or loss of the proteins NPC2 and NPC1, which are necessary to harness cholesterol from the lysosomal lumen and transfer it to the ER membrane, respectively (40–42).

To test the hypothesis that cholesterol transfer from lysosomes to the ER was involved in the transcriptional repression of cholesterol synthesis enzymes in respiratory chain (RC)-deficient cells, we treated NDUFS4-KO and WT fibroblasts with the NPC1 inhibitor U18666A (10µM, 4h), thus blocking a key protein involved in lysosome-ER cholesterol transfer. We observed that inhibition of NPC1 is sufficient to abolish the transcriptional repression of cholesterol synthesis enzymes in NDUFS4-KO cells (Figure 5A). Notably, the treatment with U18666A resulted in similar induction of transcripts of cholesterol enzymes in NDUFS4-KO and WT cells (**Figure 5A**).

**Figure 5.**
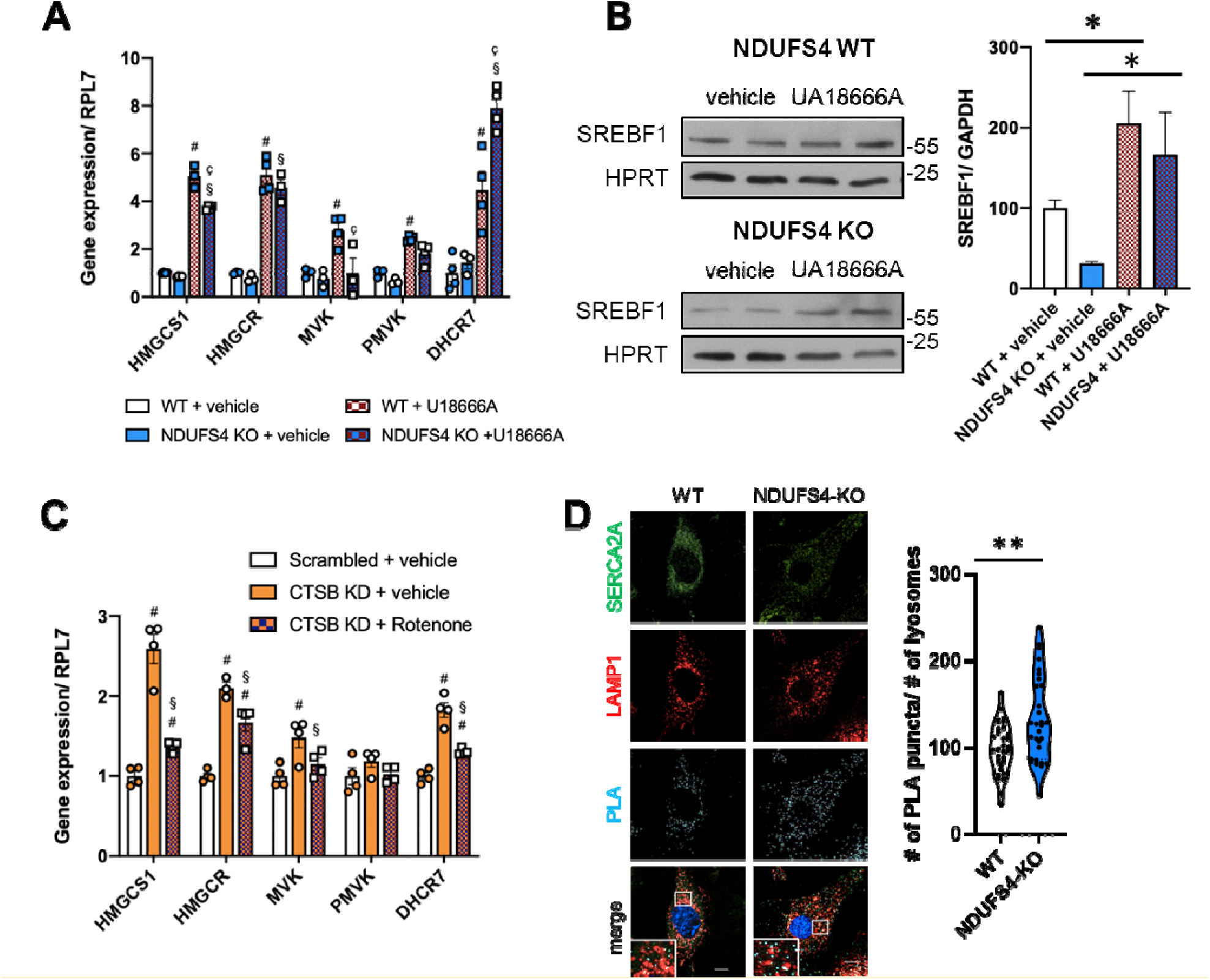
The mechanisms of cholesterol trafficking and homeostasis are fully functional in RC-deficient cells. **(A)** RT qPCR showing the transcript level of cholesterol synthesis enzymes in WT and NDUFS4 KO MEF cells untreated and treated with the NPC1 inhibitor U18666A (4h, 10µM). At least 3 independent samples were evaluated for the experiments. Two-way ANOVA with Tukey’s multiple comparisons test was used to determine the statistical significance. #= significant difference from control, §= significant difference from NDUFS4 KO, Ç= significant difference from control + U18666A. **(B)** Western blot analysis showing SREBF1 level in WT and NDUFS4 KO MEFs untreated and treated with U18666A (4h, 10µM). HPRT was used as the loading control. * p<0.05 t-test **(C)** RT qPCR of genes involved in the cholesterol synthesis performed in control and CTSB KD HeLa cells untreated and treated with the mitochondrial complex I inhibitor Rotenone (250nM, 24 h). The statistical significance was determined by two-way ANOVA with Tukey’s multiple comparisons test, evaluating at least 3 independent samples, #= significant difference from control, §= significant difference from CTSB KD. **(D)** Representative confocal images showing the contact sites between the ER (detected by the marker SERCA2A) and Lysosomes (marked with LAMP1) evaluated by proximity ligation (PLA) assay. Right: quantification of the number of PLA puncta counted on the area occupied by lysosomes. At least 30 cells were analyzed. Scale bar: 10 μm. Statistical significance was evaluated using unpaired t-test (** p < 0.01).

The profound effect of the NPC1 inhibitor suggests that the transfer of cholesterol from lysosomes to the ER occurs undisturbed in NDUFS4-KO cells. Furthermore, inhibition of NPC1 results in decreased ER cholesterol, and therefore this result further demonstrates that the cholesterol homeostatic mechanisms in the ER are functional in these cells. To confirm this, we checked the protein levels of mature SREBF1, and observed that they were significantly increased, both in WT and NDUFS4-KO, upon NPC1 inhibition (**Figure 5B**). It is noteworthy that the protein levels of NPC1 and NPC2, two key enzymes involved in lysosomal cholesterol efflux towards ER, are not changed in NDUFS4-KO cells compared to WT (**Supplementary Figure 3A**).

To further test that depletion of ER cholesterol levels could increase the expression of cholesterol synthesis enzymes in RC-deficient cells, we added 2-hydroxypropyl-beta-cyclodextrin (a cholesterol-depleting metabolite) to the WT and NDUFS4-KO cells, and observed that both WT and NDUFS4-KO show a robust increase in the transcript levels of cholesterol synthesis enzymes when cholesterol is depleted (**Supplementary Figure 3B**). Notably, this effect was also observed in cells with lysosomal defects (CTSB-KO, CTSL-KO), showing that these cells are also able to respond dynamically to changes in cholesterol levels (**Supplementary Figure 3C**).

Together, these results underscore that RC-deficient cells can dynamically respond to changes in ER cholesterol content, but that in these cells and under basal conditions there seems to be increased cholesterol egress from lysosomes to ER, thus decreasing the activity of SREBF1 and the transcript levels of its target genes.

As shown above, the response of cholesterol homeostasis to lysosomal defects is modified in cells which have impairments in the mitochondrial respiratory chain. To further test this premise using a pharmacological approach, we sought to inhibit the respiratory chain in cells with lysosomal defects which have increased expression of cholesterol synthesis enzymes. Thus, we treated HeLa CTSB-kd cells with the respiratory chain complex I inhibitor rotenone (250nM for 24h), and observed a robust decrease of the transcript levels of cholesterol synthesis enzymes (**Figure 5C**). This result supports the concept that transcriptional up-regulation of cholesterol synthesis in response to lysosomal defects requires a functional respiratory chain.

The distribution of cholesterol through cellular membranes relies on organelle contact sites. In cells lacking NPC1, there are less contacts between lysosomes and ER and thus less delivery of cholesterol from the lysosomes to the ER, which is a trigger for increased cholesterol synthesis (44). Therefore, we tested if the lysosome-ER contacts were affected in NDUFS4-KO cells. We used a proximity-ligation (PLA) assay that emits a fluorescent signal when the antibodies against the lysosomal marker LAMP1 and the ER marker SERCA2A are closer than 40nm, and therefore within the expected range of membrane contact sites (45). With this LAMP1-SERCA2A PLA assay, we observed an increase in lysosome-ER contacts in NDUFS4-KO cells (**Figure 5D**). Notably, the contacts between mitochondria-lysosomes were not affected in NDUFS4-KO cells (**Supplementary Figure 3E**). These results suggest that the increased association of lysosomes and ER in cells with RC-deficiency may contribute to enhanced lysosomal cholesterol delivery to the ER membrane leading to down-regulation of cholesterol synthesis in RC-deficient cells.

### The down-regulation of cholesterol synthesis pathway in cells with mitochondrial malfunction is independent of AMPK signaling

AMPK responds to acute mitochondrial stresses, and represses the activity of HMGCR (46) and through it the metabolic flux through the cholesterol synthesis pathway. Therefore, we sought to test whether AMPK signaling was involved in the down-regulation of expression of cholesterol synthesis enzymes in RC-deficient cells. We already showed that the HeLa UQCRC1-kd cells have decreased AMPK signaling (7). Thus, we tested if ablating AMPK activity would impact cholesterol synthesis in respiratory chain deficient cells. For that, we used MEFs lacking both α1 and α2 AMPK catalytic subunits (AMPK double knockout, henceforth AMPK-DKO), and thus devoid of AMPK activity. It is noteworthy to that the AMPK-DKO cells also show impaired lysosomal function, with lysosomal swelling and Ca2+ accumulation and decreased lysosomal hydrolysis (7). The transcript levels of cholesterol synthesis enzymes are globally up-regulated in AMPK-DKO cells (**Figure 6A**). To test if these cells are able to dynamically respond to perturbations in cholesterol homeostasis, we treated them with the NPC1 inhibitor U18666A (10µM, 24h), and observed an increase in the expression of cholesterol synthesis enzymes both in WT and AMPK-DKO cells (**Supplementary Figure 4A**). The levels of mature SREBF1 are increased in DKO cells, in agreement with the higher transcript levels of SREBF1 targets (**Figure 6B**). No change was observed in AMPK activation in WT cells treated with U18666A (**Supplementary Figure 4B**). Therefore, lower AMPK activity cannot explain the decrease in the transcript levels of cholesterol synthesis enzymes observed in respiratory chain-deficient cells and tissues. Furthermore, reactivation of AMPK in UQCRC1-kd cells was not sufficient to restore the transcript levels of cholesterol synthesis enzymes to control levels (**Supplementary Figure 4C**).

**Figure 6.**
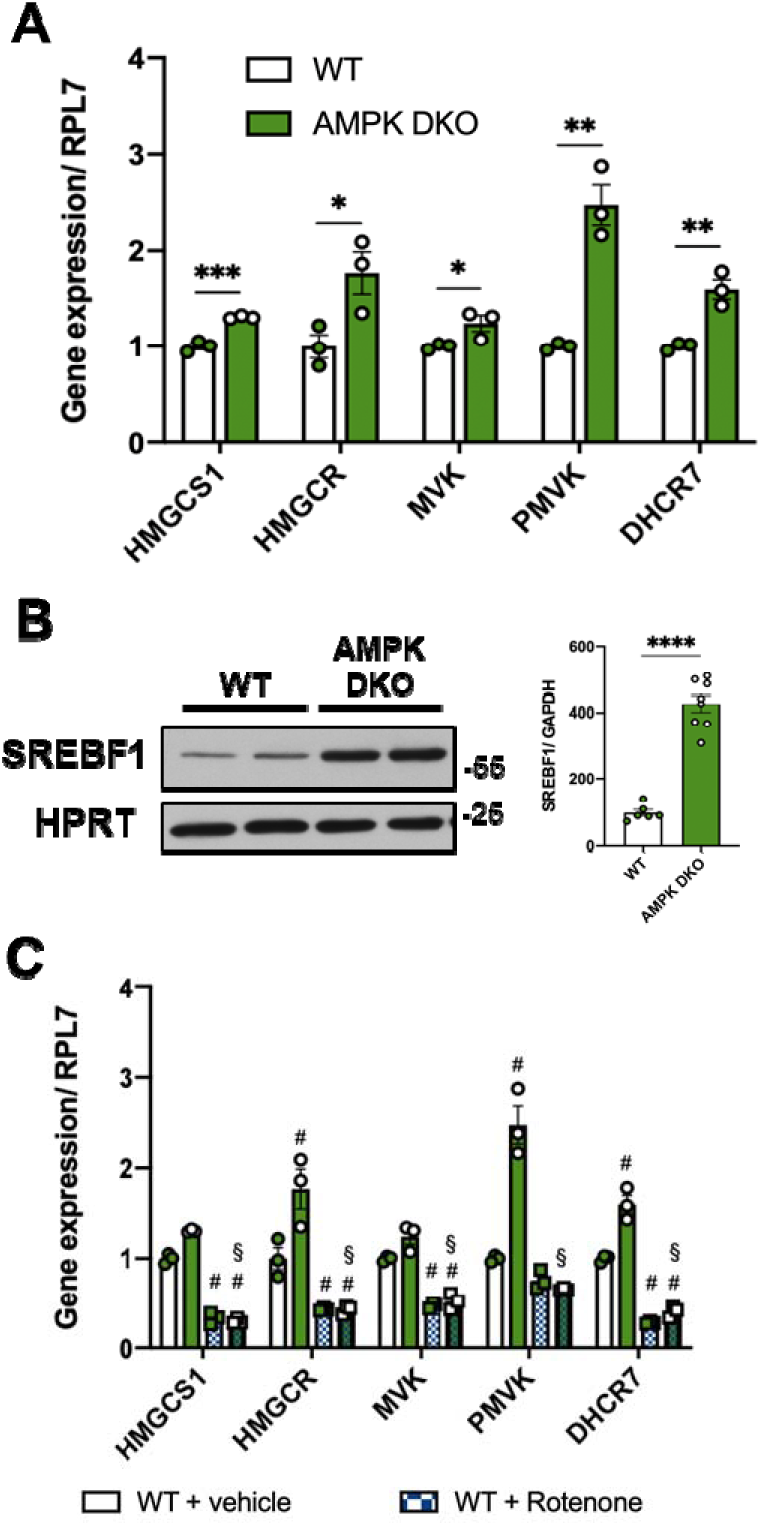
Downregulation of cholesterol synthesis in cells with mitochondrial dysfunction is independent of AMPK. **(A)** RT qPCR of cholesterol synthesis genes in control and AMPK DKO MEFs. 3 independent samples were evaluated. Statistical significance between control and mutant samples for each gene was determined by unpaired t-test, * p < 0.05, ** p < 0.01, *** p < 0.001. **(B)** Western blot analysis showing the level of SREBF1 in WT and AMPK DKO MEFs. At least 6 independent samples were analyzed using unpaired t-test, **** p < 0.0001. **(C)** RT qPCR performed in WT and AMPK DKO MEFs untreated and treated with Rotenone. 3 independent samples were evaluated. The statistical analysis was carried on using the two-way ANOVA with Tukey’s multiple comparisons test, #= significant difference from control, §= significant difference from AMPK DKO.

To verify that the response of cholesterol synthesis regulation to acute inhibition of the respiratory chain was independent of AMPK, we treated AMPK-WT and AMPK-DKO cells with the RC complex I inhibitor rotenone, and observed a global down-regulation of the transcripts of cholesterol synthesis enzymes both in WT and AMPK-DKO cells (**Figure 6C**).

AMPK and mTORC1 cross-regulate each other. Furthermore, mTORC1 also regulates cholesterol synthesis, and has previously been shown to be hyperactive in the brain of NDUFS4-KO mice (12). To test if mTORC1 was involved in the transcriptional regulation of cholesterol synthesis in RC-deficient cells, we treated NDUFS4-KO cells with torin, an established inhibitor of mTORC1 (47). While Torin treatment effectively blocked mTORC1 in both WT and NDUFS4-KO cells (**Supplementary Figure 4D**) and lowered the transcript levels of cholesterol enzymes in WT cells (**Supplementary Figure 4E**), there was no effect of mTORC1 inhibition on the transcript levels of cholesterol enzymes in NDUFS4-KO cells (**Supplementary Figure 4E**). These results suggest that mTORC1 is not involved in the down-regulation of the transcript levels of cholesterol synthesis enzymes in NDUFS4-KO cells.

### MCOLN1 loss-of-function decreases cholesterol synthesis in cells with mitochondrial defects

The observation that mitochondrial defects result in decreased expression of cholesterol synthesis enzymes, while lysosomal defects have the opposite effect, implies that the changes in the expression of the cholesterol pathway in mitochondrial defects cannot be explained simply by decreased lysosomal hydrolysis, as demonstrated above. Furthermore, changes in AMPK and mTORC1 activity also cannot explain this differential effect. We have previously shown that in cells with respiratory chain deficiency such as HeLa UQCRC1-kd and NDUFS4-KO mouse fibroblasts, there is decreased activity of TRPML1 (also known as mucolipin 1, MCOLN1), a lysosomal export channel for divalent cations particularly implicated in lysosomal Ca^2+^ efflux (48). Thus, we tested if loss of MCOLN1 activity impacts the expression of cholesterol synthesis enzymes. We used mouse fibroblasts lacking MCOLN1 (MCOLN1-KO) and the corresponding WT cells, and measured the transcript levels of cholesterol synthesis enzymes by qPCR. We observed that the transcript levels of the cholesterol synthesis enzymes are down-regulated in MCOLN1-KO (**Figure 7A**). This result is similar to what is observed in cells with mitochondrial defects, suggesting that the loss of MCOLN1 activity, and corresponding accumulation of Ca^2+^ and possibly other divalent ions in the lysosome, correlates with the repression of the cholesterol synthesis pathway. To further demonstrate that the decrease in MCOLN1 activity was causative of the decreased transcript levels of the cholesterol synthesis enzymes, we tested if reactivating the MCOLN1 channel in respiratory chain-deficient cells would affect the expression of cholesterol synthesis enzymes. We treated the NDUFS4-KO cells with the MCOLN1 activator MLSA1, and observed an increase in the transcript levels of cholesterol synthesis enzymes (**Figure 7B**). We obtained a similar result when treating UQCRC1-kd cells with MLSA1 (**Figure 7C**), suggesting that lower MCOLN1 activity in cells with defective respiratory chain is a key mediator of cholesterol synthesis repression. These results highlight that MCOLN1 activity impacts the regulation of cholesterol homeostasis, akin to other roles played by this protein in nutrient sensing (49).

**Figure 7:**
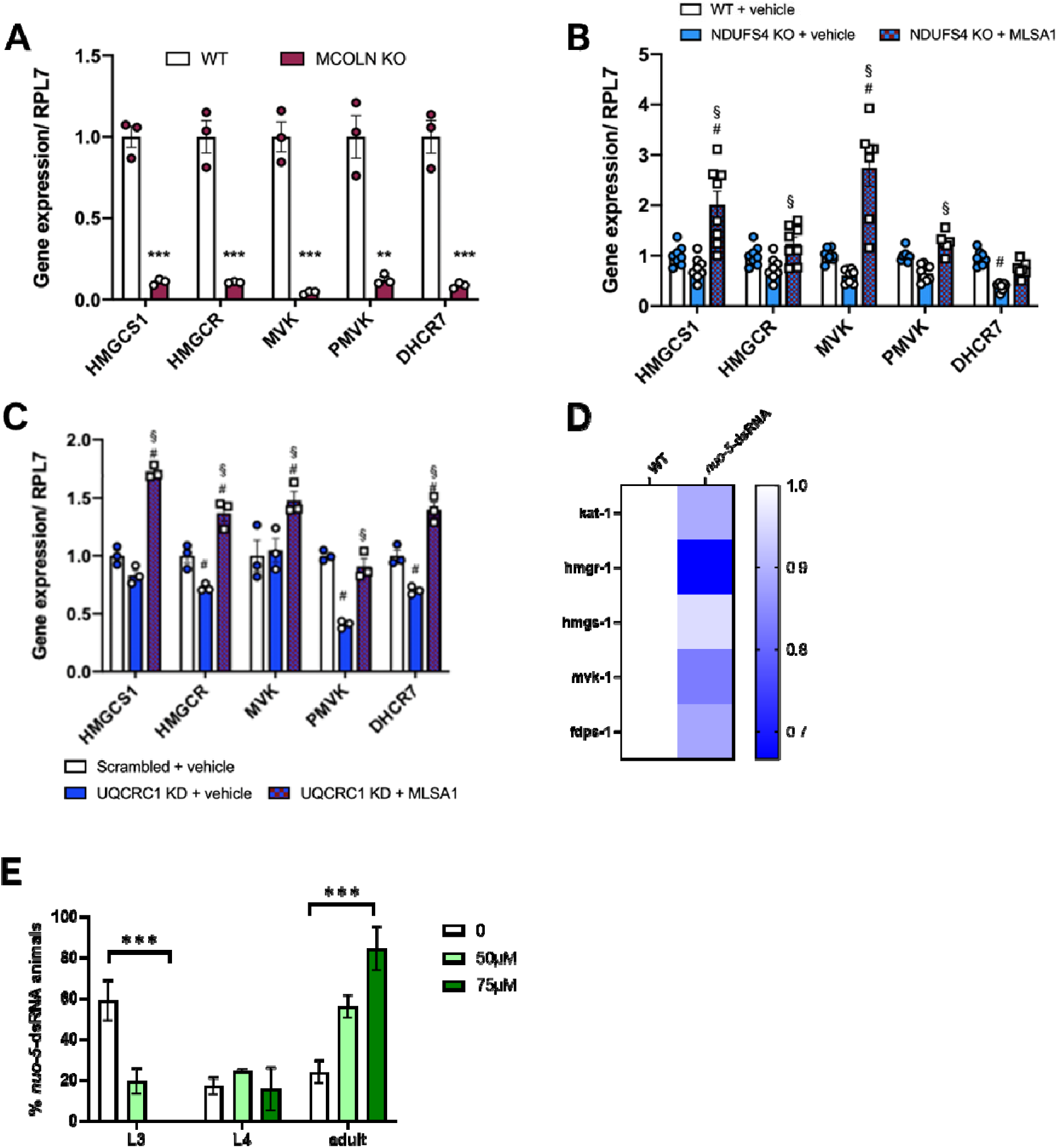
MCOLN1 loss-of-function decreases cholesterol synthesis in cells mitochondrial deficiency cells and improves an *in vivo* model of mitochondrial disease. **(A)** RT qPCR of genes involved in the cholesterol biosynthesis pathway in WT and MCLN KO MEFs. 3 independent samples were evaluated. The statistical significance was determined by two-way ANOVA with Tukey’s multiple comparisons test. ** p < 0.01, *** p < 0.001. The same genes were analyzed through RT qPCR in WT and NDUFS4 KO MEFs untreated and treated with the MCOLN1 activator MLSA1 **(B)**. At least 4 independent replicates were evaluated. Statistical significance was evaluated with two-way ANOVA with Tukey’s multiple comparisons test. #= significant difference from control, §= significant difference from NDUFS4 KO. RT qPCR of these genes was also performed in control and UQCRC1 KD Hela cells, untreated and treated with MLSA1 **(C)**. 3 independent samples were used and analyzed using two-way ANOVA with Tukey’s multiple comparisons test. #= significant difference from scrambled, §= significant difference from UQCRC1 KD. **(D)** Heatmap showing the fold change in the transcript levels of the indicated genes in nuo-5 worms relative to their parental control line. **(E)** Percentage of *nuo-5*-dsRNA worms in larval 3 (L3), larval 4 (L4) and adult stages, in regular NGM in the absence or presence of MLSA1 (50μM or 75μM), after 4 days, from eggs laid from wild-type animals fed bacteria transformed either with empty vector (pl4440) or with dsRNA against C. elegans complex I subunit NDUFS1 (*nuo-5*). ***p<0.0001 two-way ANOVA.

To test if activation of MCOLN1 might have benefits in vivo, we took advantage of a C. *elegans* model of mitochondrial respiratory chain deficiency due to the silencing of complex I subunit *nuo*-5 (an orthologue of mammalian complex I subunit NDUFS1), which we previously characterized (50). The *nuo-5*-dsRNA worms show larval arrest primarily in the L3 stages, and a delay in reaching adulthood (only about 25% of *nuo-5*-dsRNA worms reach adulthood on day 4, compared to 100% of WT worms who reach adulthood on day 3) (50). While C. *elegans* do not synthesize cholesterol, the mammalian cholesterol pathway is conserved in worms until the synthesis of mevalonate (51). Of note, because worms lack the ability to synthesize cholesterol from mevalonate, they rely on exogenous cholesterol, and decrease in cholesterol availability in the diet results, similar to mitochondrial deficiency, in developmental arrest (52). It is thus possible that the decrease in the expression of mevalonate synthesis enzymes in *nuo-5*-depleted worms contributes to lower availability of mevalonate and to the developmental delay. In support of this possibility, we used a transcriptome dataset of *nuo-5*-dsRNA worms that we previously published (50) to assess the expression of mevalonate synthesis enzymes, and observed a global down-regulation across the pathway (**Figure 7D**; the transcripts for PMVK, MVD and IDI1 were not detected in the dataset). Interestingly, we previously identified an acetylcholine signaling defect in the *C. elegans* mitochondrial disease model (50), a cellular function which is tightly linked to cholesterol homeostasis (53). Given the results obtained in cell lines with reactivation of the lysosomal channel MCOLN1, we thus sought to test if treating the worms with the MLSA1 would be beneficial. We titrated MLSA1 addition to the culture medium of *nuo-5*-dsRNA, in two different doses, 50µM and 75µM, and observed that the percentage of worms reaching adulthood by day 4 increased proportionally to the concentration of MLSA1 (**Figure 7E**). While the consequences of MLSA1-induced MCOLN1 reactivation are likely broad and go beyond the effect on the expression of cholesterol/mevalonate synthesis enzymes, these results show that the pathological consequences triggered by defects in the respiratory chain can be, at least in part, rescued by manipulating the lysosome which, among other pathways, will improve cholesterol homeostasis.

## DISCUSSION

We show here that the transcriptional regulation of the cholesterol synthesis pathway is affected both by mitochondrial defects and by lysosomal defects, in an opposite manner. These data are supported by whole cellular transcriptome of different mitochondrial and lysosomal defects, as well as tissues from mouse models of mitochondrial or lysosomal defects.

Cholesterol is an important constituent of cellular membranes, and therefore its levels are subject to a complex regulatory network that involves multiple cellular organelles and proteins to regulate cholesterol synthesis at transcriptional level as well as post-translationally. At the transcriptional level, the regulation of cholesterol synthesis relies on the SREBPs transcription factors, of which SREBF1-a and SREBF2 are the predominant isoforms of SREBPs in cultured (mammalian) cell lines (37). Here, we observed an increase in the transcript levels of cholesterol synthesis enzymes in multiple cell models with different types of lysosomal defects, as well as in mouse models of lysosomal deficiency. Conversely, in cellular and mouse models of mitochondrial respiratory chain deficiency, we observed a decrease in the transcript levels of cholesterol synthesis enzymes. We also observed a decrease in the transcript levels of mevalonate synthesis enzymes in worms silenced for *nuo-5*, a subunit of respiratory chain complex I, indicating that the effect is conserved even in nematodes. While SREBF1 is usually seen as a generic activator of lipid synthesis and SREBF2 a more specific activator of the cholesterol synthesis pathway (54), here we observed that, in the cellular models of mitochondrial or lysosomal defects, the changes in the transcript levels of the cholesterol synthesis enzymes were paralleled by changes in SREBF1 protein levels and intracellular localization, while SREBF2 levels remained unchanged. In cellular models of lysosomal defects, SREBF1 was more abundant in the nucleus, in agreement with the increased transcript levels of the cholesterol synthesis enzymes. In cells with mitochondrial respiratory chain defects, there was less SREBF1 protein, and lower transcript levels of the same enzymes. Because the transcript levels of SREBF1 were unchanged in the cells with lysosomal or mitochondrial defects, a decrease in the protein levels of SREBF1 in cells with mitochondrial defects could be due to proteolytic degradation. Interestingly, we observed that the decrease in SREBF1 protein levels could be prevented by inhibition of the proteasome, supporting the notion that SREBF1 was being actively targeted for degradation in the cellular models of respiratory chain deficiency. It has been previously described that SREBF1 can be subjected to ubiquitin-mediated proteolysis, and that the phosphorylation of SREBF1 by GSK3B promotes this process (38). Accordingly, we found that GSK3B was hyperactive in the cellular models of respiratory chain deficiency, providing a route to explain why SREBF1 levels were decreased in those cells. Activation of GSK3B has been reported in cells with 1-methyl-4-phenylpyridinium iodide (MPP(+))-induced mitochondrial malfunction (55). A study of urinary stem cells from patients of MELAS (Mitochondrial Encephalopathy, Lactic Acidosis and Stroke-Like Episodes) shows a decrease in GSK3B phosphorylation, which would correlate with increased activity, but the total levels of GSK3B were not measured and therefore it is not clear if the protein is indeed active (56). In contrast, human fibroblasts with mutations in NPC1, which result in lysosomal cholesterol storage, show an increase in the inhibitory phosphorylation on GSK3B, and therefore decreased GSK3B activity (57). While this result needs to be tested in a more systematic manner, it is possible that the differential activation of GSK3B by mitochondrial and lysosomal defects contributes to the opposite regulation exerted over the transcription of the genes encoding cholesterol synthesis enzymes. The mechanism that regulates GSK3b in response to defects in mitochondria or in lysosomes warrants further investigation. It is also possible that, in addition to cholesterol, other lipids are affected, given the lipogenic role of SREBF1.

The up-regulation of cholesterol synthesis at transcript level by different lysosomal defects has been shown. Some of these defects include absence or loss-of-function of the proteins NPC1 and NPC2, which are involved in the lysosomal egress of cholesterol and sphingomyelin (17). Other lysosomal defects or interventions that impact acidification also result in transcriptional activation of cholesterol synthesis (27, 58, 59). The models of lysosomal defects that we assessed are knock-out of four different lysosomal genes: GAA, involved in lysosomal glycogen hydrolysis (60), cathepsins B and L, which are involved in lysosomal proteolysis (61), and LAMP2, involved in chaperone-mediated autophagy and cholesterol trafficking (62). All these mouse fibroblasts with different lysosomal defects yielded similar consequences to the transcriptional regulation of cholesterol synthesis. These results were also similar to those we obtained in HeLa cells with lysosomal defects, indicating that the effect is likely valid across cell types. Furthermore, we obtained similar results from the brain and liver of the mouse models studied, indicating a conserved effect across tissues. It is important to note that the transcriptional regulation of cholesterol synthesis in the liver is not only dependent on SREBPs, and that nuclear receptors such as liver X receptors (LXRs) may also be involved (63).

The lysosomes are intracellular logistics platforms, processing and sensing several nutrients and distributing them to other cellular regions. They communicate with other organelles such as the endoplasmic reticulum (ER) and mitochondria. It is known that perturbation of lysosomal contact sites with the ER results in lysosomal cholesterol storage and increased association between lysosomes and mitochondria (44). In the cellular model of respiratory chain deficiency, we did not observe any difference in the number of mitochondria-lysosome contact sites, but the contacts between the lysosomes and the ER were increased, which may facilitate transfer of cholesterol between the ER and the lysosomes. Cholesterol transfer at ER-lysosome contact sites can occur in both directions (64), but given that there is less SREBF1 in the models of mitochondrial malfunction, it is unlikely that there is net transfer of cholesterol from ER to lysosomes in these cells. The contribution of ER-lysosome-mitochondria cholesterol transfer through contact sites to the transcriptional regulation of cholesterol in models of mitochondrial and lysosomal malfunction remains to be determined.

We show here that the effect of lysosomal malfunction on cholesterol synthesis requires a functional respiratory chain. When we perturb mitochondrial respiration in cellular models of lysosomal deficiency, the transcriptional up-regulation of cholesterol synthesis subsides. This observation raises tantalizing possibilities of targeting the respiratory chain to lower cholesterol synthesis in models of hypercholesterolemia. For example, metformin, a respiratory chain complex I inhibitor that is widely used as a drug against type II diabetes, reduces serum cholesterol levels in patients (65, 66). Cells treated with metformin also show a transcriptional down-regulation of cholesterol synthesis (67, 68), although this may involve metformin-induced AMPK activation (69). In our cellular models of mitochondrial malfunction, we did not observe a role for AMPK or mTORC1 in the transcriptional regulation of cholesterol synthesis. Notably, acute pharmacological inhibition of different respiratory chain complexes was also shown to have an inhibitory effect on the transcription of cholesterol synthesis enzymes in cells (70, 71).

We showed previously that lysosomes accumulate Ca^2+^ in models of mitochondrial respiratory chain deficiency, due to decreased activity of the lysosomal Ca2+-export channel MCOLN1 (7). Interestingly, we observed that treatment of respiratory chain-deficient cells with an MCOLN1 agonist normalized the transcript levels of cholesterol synthesis enzymes. Furthermore, we observed that pharmacological activation of MCOLN1 significantly rescued the developmental delay that characterizes the worms silenced for *nuo-5*, a model of mitochondrial respiratory chain deficiency, suggesting that the normalization of cholesterol homeostasis contributed to the systemic improvement in the *nuo-5*-treated worms. Future studies will further detail the mechanisms by which mitochondrial and lysosomal defects impact the transcriptional regulation of cholesterol synthesis, and the physiopathological impacts of this process.

## METHODS

### Cell culture and treatments

Perinatal mouse fibroblasts were cultured in Dulbecco’s modified Eagle’s medium (DMEM, Gibco) supplemented with 10% v/v fetal bovine serum (FBS) and 1% Penicillin/Streptomycin (Corning). Cell lines were maintained at 37°C in a 5% CO2 controlled atmosphere. TrypLE (Gibco) was employed to detach cells and generate subcultures. When at 80% of confluency, cells were treated with, MG132 (10uM, 2h), U18666A (10µM, 24h), with MLSA1 (20 uM, 4h). The NDUFS4-KO cells and the GAA-KO cells were obtained as described earlier (7, 18). The CTSB-KO, CTSL-KO mouse embryonic fibroblasts were a kind gift from Prof. Thomas Reinheckel (Albert-Ludwigs-Universität Freiburg, Germany). The LAMP2-KO mouse embryonic fibroblasts were a kind gift from Prof. Paul Saftig (University of Kiel, Germany). The AMPKα1α2-double knockout mouse embryonic fibroblasts were a kind gift from Prof. Benoit Violet (Institut Cochin, Paris).

### Generation of stable knockdown cells

Lentiviral stable knockdown generation was done by growing HEK293T packaging cells in DMEM high glucose supplemented with 10% FBS and after 24 h were transfected with viral components and shRNA against target genes (or scrambled control) using Lipofectamine 2000 (Invitrogen, 11668–019), grown and concentrated using Lenti-X Concentrator (Clontech Laboratory, 631232) according to the manufacturer’s instructions. The shRNA constructs were purchased from Open Biosystems (Dharmacon). HeLa cells were then seeded at 12000 cells/cm2 and grown overnight to 70–80% confluence. These cells were transduced with the lentiviral particles using Polybrene (8 μg/ml; Sigma-Aldrich TR-1003-G). Puromycin (Fisher Scientific, BP2956-100; 4 ug/ml) was used as selection agent. These knocked-down cells were selected with puromycin, the resistance marker of the shRNA vector, and stable silenced lines were established (Supplementary Figure 1).

### RNA sequencing

RNA sequencing was performed as described (72). RNAseq data analysis was performed using Partek Software Suite. RNAseq data was aligned to the reference genome mm10 by the BowTie algorithm, and the transcripts were quantified using the ‘mm10 - Ensembl Transcripts release 95’ as reference. For differential expression analysis, Bonferroni multi-test correction was applied, and adjusted p-value<0.05 was considered significant. Ingenuity Pathway Analysis was used for assessment of transcription factor activity, as described (27). The data is deposited at Gene Expression Omnibus (GSE256471).

### Mouse tissues

The NDUFS4-KO and GAA-KO mouse tissues were obtained as described earlier (Lorena Autophagy, King eLife1, eLife2). All animal experiments were carried out in accordance with the European guidelines for animal welfare and were approved by the Lower Saxony Landesamt fur Verbraucherschutz and Lebensmittelsicherheit (LAVES) registration number 15–883.

### C. *elegans* strains and maintenance

We employed standard nematode culture conditions. All strains were maintained at 20 °C on Nematode Growth Media (NGM) agar supplemented with Escherichia coli (OP50 or transformed HT115), unless otherwise indicated. The dsRNA transformed bacteria for feeding were derived from the Ahringer C. elegans RNAi library (73): nuo-5 (Y45G12B.1).

### Development assay and treatment with MLSA1

Both dsRNA bacterial clones (sequence validated), were grown in LB media to a concentration of 0.9 OD and spotted on NGM plates. Prior the addition of MLSA1, dried bacteria were inactivated by UV light exposure for 25 min. Afterward, the compound dissolved in DMSO was spotted at the desired final concentration (namely 50µM and 75µM). Plates were allowed to dry overnight at room temperature and subsequently stored at 4°C. The first generation of nematodes (P0) was grown on RNAi plates without the compound. The second generation (F1) was grown on the MLSA1-supplemented plates. The developmental stage of the F1 was checked on the second, third and fourth day after egg-lay. The percentage of animals in each developmental stage was calculated at each time point, and the data were pooled from two independent experiments.

### Cell lysis and immunoblotting

Cells were harvested in lysis buffer (N-dodecylmaltoside (Dot Scientific) 1,5% w/v dissolved in phosphate buffered saline (PBS) supplemented with protease and phosphatase inhibitor cocktail (Thermo Scientific). Lysates were incubated on ice for 30 minutes and cleared by centrifugation at 16000 x g for 30 minutes at 4°C. Supernatants were collected and stored at -20 C° until used. Protein concentration in the samples was evaluated using the Pierce BCA Protein Assay Kit (Thermo Scientific) following the manufacturer’s instructions. 50 ug of protein lysates were boiled for 10 minutes at 100 C° and loaded 13% Tris-glycine polyacrylamide gels in SDS/Tris-glycine running buffer. PageRuler plus pre-stained protein ladder (Thermo Scientific) were used for protein size estimation. The resolved proteins were transferred onto nitrocellulose membranes (Amersham) through a Trans-Blot semi-dry transfer cell (Bio-Rad). Membranes were subsequently blocked in Tris-buffered saline plus 0.1% Tween (TBS-T) plus 5% non-fat milk for 1 h at room temperature (RT) and then incubated over-night at 4 °C with primary antibodies diluted in TBS-T plus 5% non-fat milk. Membranes were then washed in TBS-T (3x10 minutes) at RT and subsequently incubated for 1h at RT with horseradish peroxidase (HRP)-conjugated secondary antibodies. Membranes were then washed in TBS-T (3x10 min) at RT and rinsed in TBS-T. Immunoreactive proteins were visualized using Immobilon® Forte Western HRP Substrate (Merck Millipore) at the Imager Fluor-Chem M (Proteinsimple). Densitometric analysis was carried out using the Image J software.

The following antibodies were used for western blotting: mouse SREBP1 (Abcam AB3259), mouse SREBP2 (RD system MAB7119), mouse GAPDH (Sigma-Aldrich G8795) rabbit HPRT (Abcam, AB10479), rabbit Insig1 (Abcam,AB70784), rabbit SCPAP (Abcam, AB153933), rabbit PARP (cell signaling technology, 542S), rabbit GSK3-beta (Cell signaling technology D5C5Z), rabbit SGK3-alpha/beta phosphor S21/S9 (Cell signaling technology, D8SeE12), PDI (Cell signaling technology, C81H6).

### Cell fractionation

Cell fractionation of nuclear and cytoplasmic compartments was performed with the NE-PERTM Nuclear and Cytoplasmic Extraction Reagents (Thermo-Scientific) kit following the manufacturer’s instructions.

To obtain ER fractions, NDUFS4 WT and KO MEF cells grown on 15-cm dishes were resuspended in 10 mM NaCl, 1,5 mM MgCl2, 10 mM Tris-HCl, pH 7,5 buffer and homogenized using an overhead stirrer (Wheaton instruments) with a Teflon pestle. Following homogenization cells were subjected to differential centrifugation to obtain nuclear, whole cell, crude mitochondria and microsomal fractions. The latter were ultracentrifuged at 100 000xg for 1h to isolate ER fractions.

### Immunocytochemistry, proximity ligation assay (PLA) and confocal imaging

Cells were cultured onto 12mm glass coverslips (Thermo-Scientific) coated with Poly-L-Lysine (Sigma-Aldrich). Cells were fixed with 4% w/v Paraformaldehyde (PFA) for 1h at RT. After cell permeabilization in PBS plus triton 0,1% for 30 minutes at RT and a blocking step performed in 3% BSA dissolved in PBS for 30 minutes at RT, cells were stained with the appropriate primary antibodies diluted in PBS plus 3% BSA overnight at 4°C. Subsequently, for immunostaining experiments cells were washed three times in PBS and incubated with the secondary antibody Alexa Fluor® 488 conjugated (Invitrogen) or Alexa Fluor® 568 conjugated (Invitrogen). To perform the PLA assay the Duolink In Situ Detection reagents (Sigma Aldrich) were used. Briefly, after primary antibodies staining, cells were incubated for 1 hour at 37°C with PLUS and MINUS PLA probes (Sigma Aldrich). Then cells were incubated with the Ligase diluted in the ligation buffer for 30 minutes at 37 °C, subsequently, the amplification step was performed by incubating cells with the polymerase diluted in the amplification buffer for 1 h at 37°C. Cells were finally incubated with Hoechst 33258 (Invitrogen) for 5 minutes at RT, and mounted with the ProLong Diamond Antifade (Invitrogen) mounting medium on glass slides.

Images were acquired with a LeicaSP7 confocal microscope (Leica Microsystems) and quantified using ImageJ.

The following antibodies were used: mouse SERCA 2ATPase (Invitrogen MA3-919), mouse HSP60, rabbit LAMP1 (Cell Signalling technology, C54H11), mouse HSP60 (Proteintech 66041-1)

### Real time PCR

RNA was extracted from MEF cells using the RNeasy Mini Kit (Qiagen) following manufacturer instructions. RNA quality and concentration were assessed by Nanodrop. cDNA was synthesized from total RNA using iScript cDNA Synthesis Kit (Bio Rad). 100 ng of cDNA have been used for each qRT-PCR reaction. RT-qPCR reaction was performed on 98-well plate adding to the cDNA the PowerUp SYBR Green Master Mix (Applied biosystem) and the DNA primers.

The following primers were used:

DHCR7. For: AGGCTGGATCTCAAGGACAAT. Rev: GCCAGACTAGCATGGCCTG HMGCR. For: TGTTCACCGGCAACAACAAGA. Rev: CCGCGTTATCGTCAGGATGA HMGCS1. For: CGGATCGTGAAGACATCAACTC. Rev: CGCCCAATGCAATCATAGGAA MVK. For: GGTGTGGTCGGAACTTCCC. Rev: CCTTGAGCGGGTTGGAGAC PMVK. For: AAAATCCGGGAAGGACTTCGT. Rev: AGAGCACAGATGTTACCTCCA. TULP3. For: CAGTGCCTTTGACGATGAGAC. Rev: TTCTGGGTTTGGCTGTACCAT. PLEKF1. For: GAGAGCTGTTTTGGGGCATC. Rev: AATACTGCCGTACACCAGGAT. IL20RB. For: ACCCCTTTAACCGAAATGCAA. Rev: CCTCCAGTAGACCACAAGGAA. SH3RF1. For: CAGGTCCATATAAGCACCACTG. Rev: GGTAGGGGACATCTGAAGGGA. NUCB2. For: GGACAAGACCAAAGTACACAACA. Rev: CCGCTCCTTATCTCCTCTATGT. SDF2L1. For: CTGCACTCACACGACATCAAA. Rev: CGCGAATCCGCCAGTAACT. SCAP: For: CCGAGCATTCCAACTGGTG. Rev: CCATGTTCGGGAAGTAGGCT. SREBP1: For:TGACCCGGCTATTCCCTGA. Rev: CTGGGCTGAGCAATACAGTTC. SREBP2: For: GCAGCAACGGGACCATTC. Rev:CCCCATGACTAAGTCCTTCAACT. RPL7. For: CTGCTGGGCCAAAAACTCTCA. Rev: CCTTCAACTCTGCGAAATTCCTT.

### Cholesterol assay

The amount of cholesterol in the cell ER fractions was evaluated through Amplex cholesterol Assay kit (Invitrogen A12216) following the manufacturer instructions.

## ACKNOWLEDGEMENTS

The conclusion of this work was supported by the start-up funds from the Department of Cellular and Molecular Physiology at Penn State University and NIH R56AG082790-01 to NR. The initial stage of this work was supported by DAAD-Germany doctoral fellowship to LP; ERC Starting Grant 337327; DFG SFB1190-P02; FCT 2022.09311.PTDC.

## AUTHOR CONTRIBUTIONS

FA, LP, JD, KFY, SM and AS performed experiments. FA, LP, KFY, NV and NR designed experiments. All authors analyzed and interpreted data. NR designed the project. FA and NR prepared the figures and wrote the manuscript, which all authors commented on.

---

**Supplementary Figure 1.**
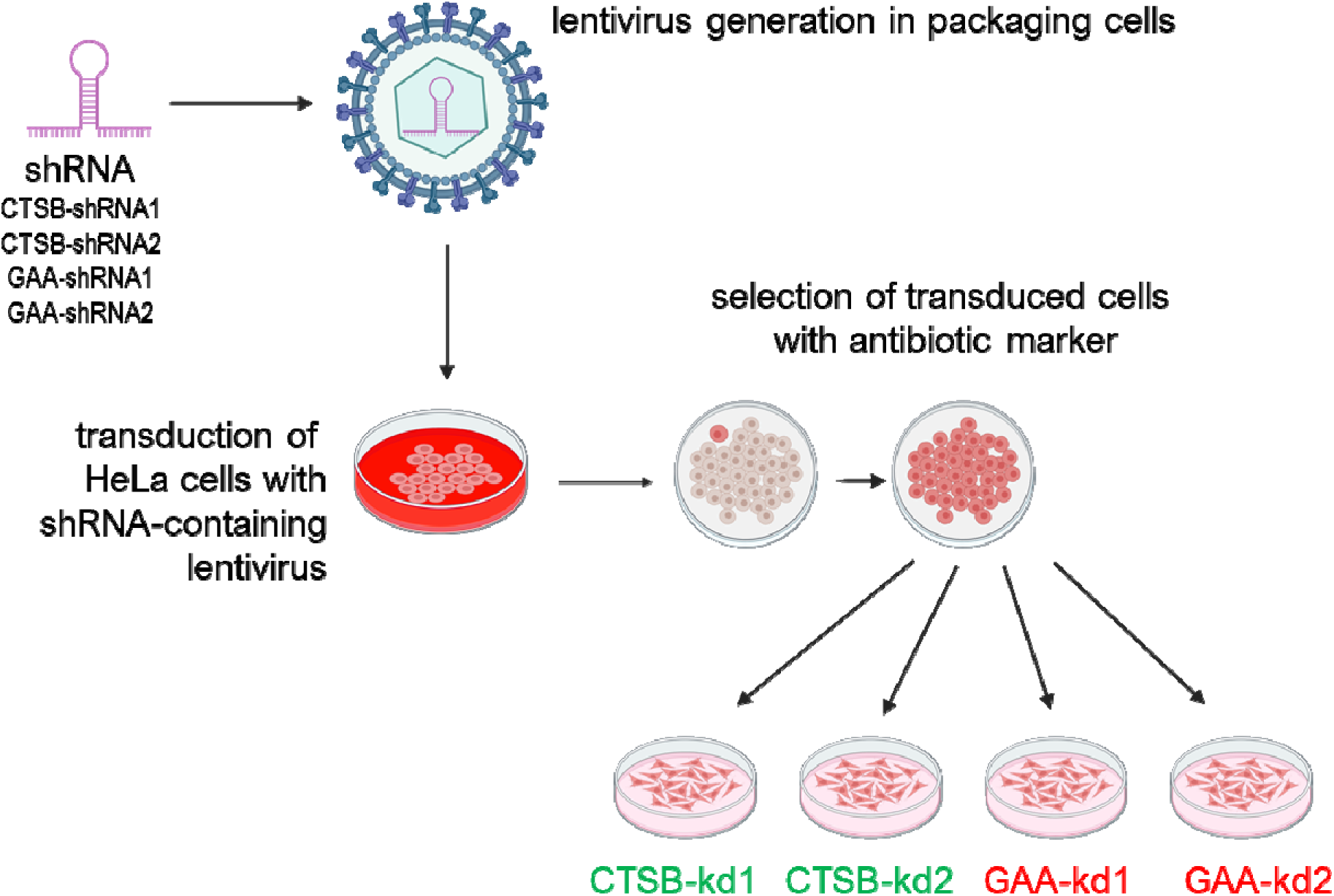
Schematic representation of the preparation of the cell lines with stable knock-downs.

**Supplementary Figure 2.**
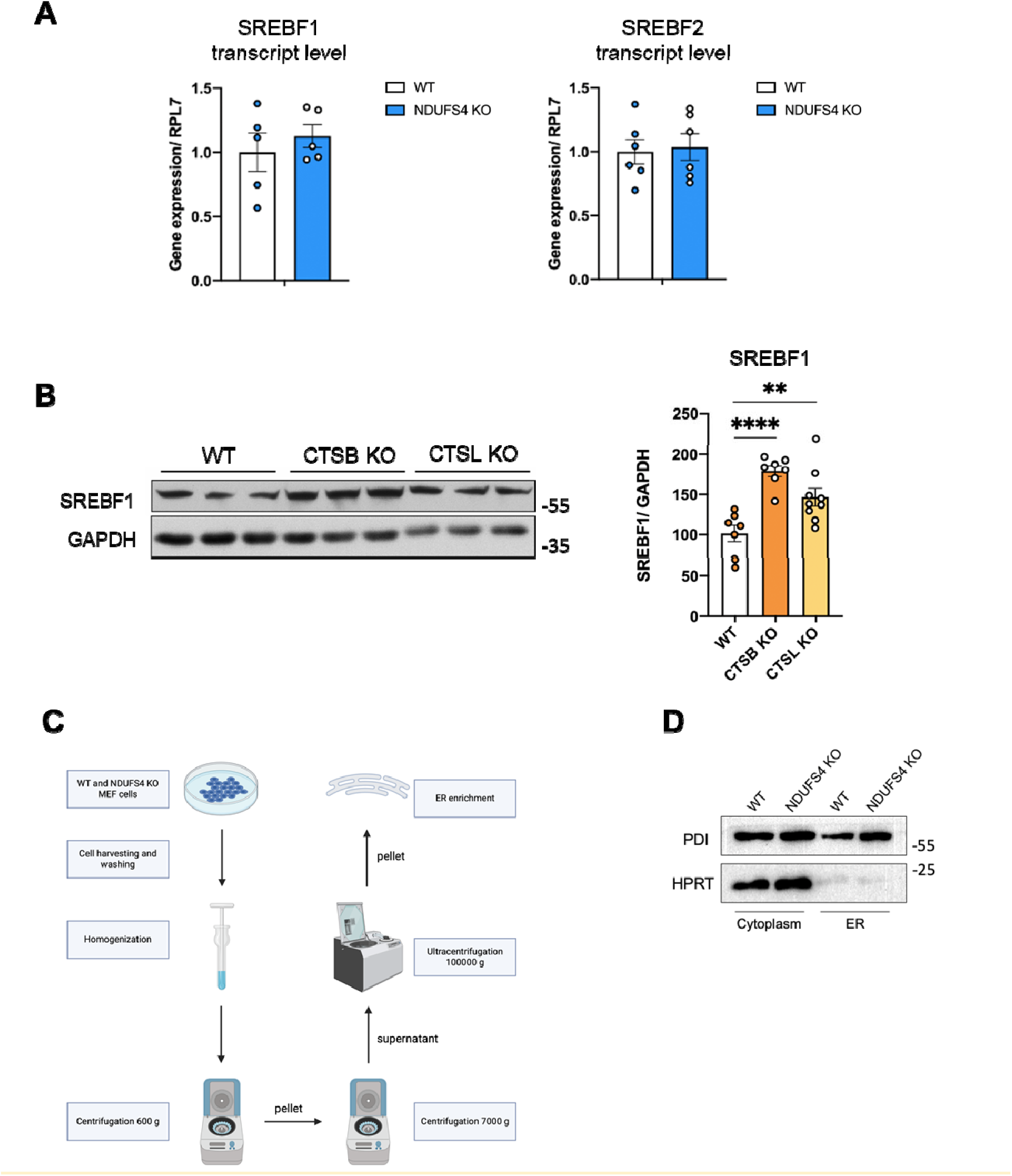
SREBF1 and SREBF2 levels are affected by mitochondrial and lysosomal deficiency. **(A)** Transcript levels of SREBF1 and SREBF2 in WT and NDUFS4-KO MEF cells, measured by qPCR. At least 6 independent samples were evaluated. Statistical significance was evaluated by unpaired t-test. **(B)** Western blot analysis of SREBF1 level in WT, CTSB KO and CTSL KO MEF cells. GAPDH was used as the loading control. At least 7 independent samples were analyzed. Statistical significance was evaluated with one-way ANOVA with Tukey’s multiple comparisons test, ** p < 0.01, **** p < 0.0001. **(C)** Schematic representation of the isolation procedure to obtain ER-enriched fractions. **(D)** Western blot confirming the isolation of the ER fraction from WT and NDUFS4-KO cells. PDI was used as ER marker, while HPRT was used as cytosolic marker. The fractions were obtained with the isolation procedure schematically represented in C.

**Supplementary Figure 3.**
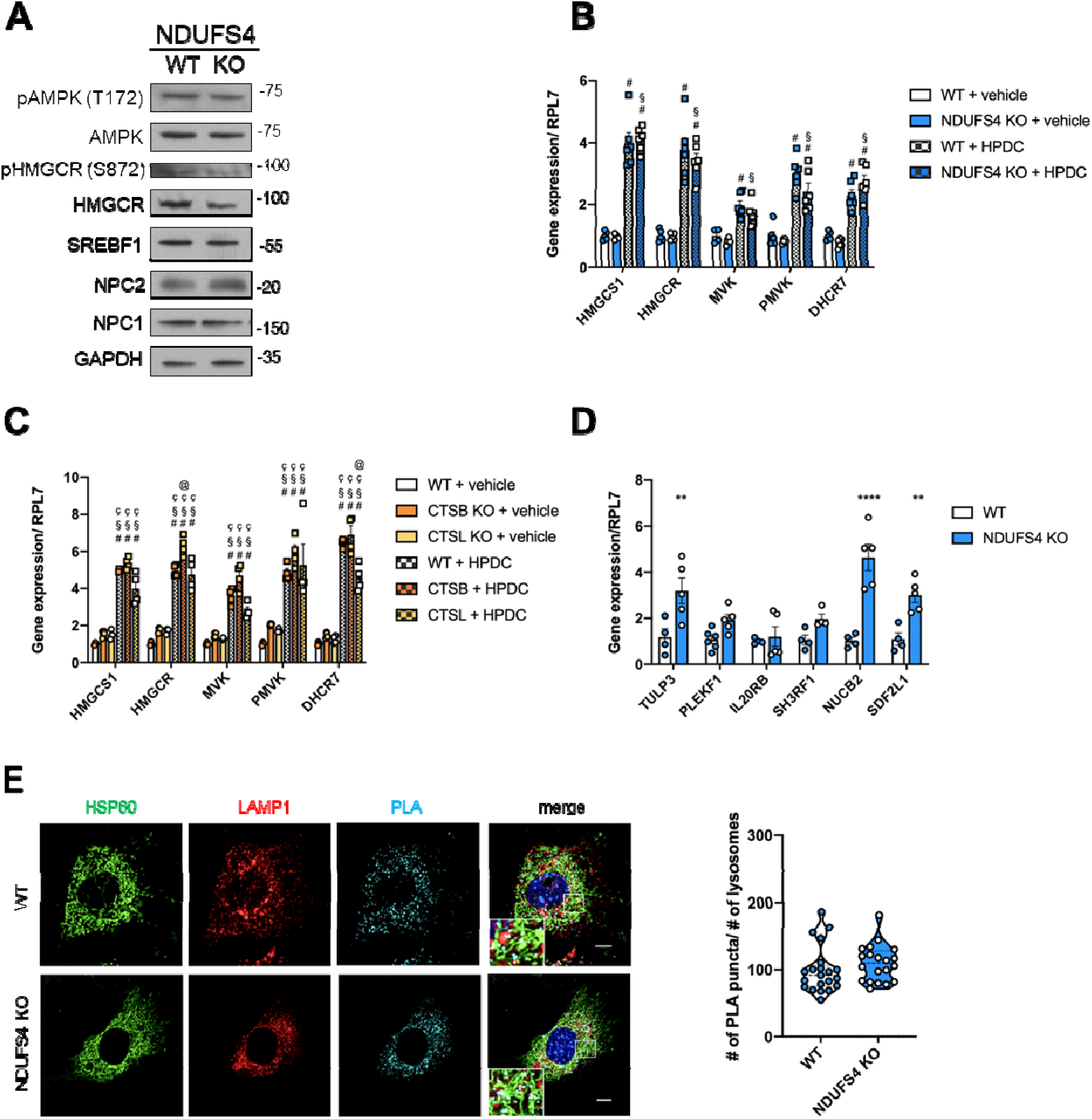
**(A)** Western blot showing the protein level of p-AMPK (Thr172), AMPK, p-HMGCR (Ser872), HMGCR, SREBF1, NPC1 and NPCT in WT and NDUFS4 KO MEF cells. GAPDH has been used as loading control. **(B)** Transcript levels of enzymes involved in the cholesterol synthesis pathway in WT and NDUFS4-KO MEF untreated and treated with the cholesterol depleting molecule HPDC (4h, 1% w/v), evaluated by qPCR. At least 3 independent samples were analyzed. The statistical significance was determined by two-way ANOVA with Tukey’s multiple comparisons test. #= significant difference from control, §= significant difference from NDUFS4 KO. The same transcripts were evaluated in control, CTSB KD and CTSL KD HeLa cells, treated or not with HPDC **(C).** Two-way ANOVA with Tukey’s multiple comparisons test was used to evaluate the statistical significance, a minimum of 3 samples were analyzed. #= significant difference from control, §= significant difference from CTSB KO, Ç= significant difference from CTSL KO, @= significant difference from control + HPDC. **(C)** Confocal imaging (scale bar: 10 μm) of mitochondria-lysosomal contact sites assessed by PLA analysis, performed using the anti-HSP60 (mitochondrial marker) and the anti-LAMP1 (Lysosomal marker) antibodies. The graph shows the number of PLA puncta counted in the area occupied by LAMP1-positive structures. At least 20 cells per genotype were analyzed. The statistical significance was evaluated with unpaired t-test.

**Supplementary Figure 4.**
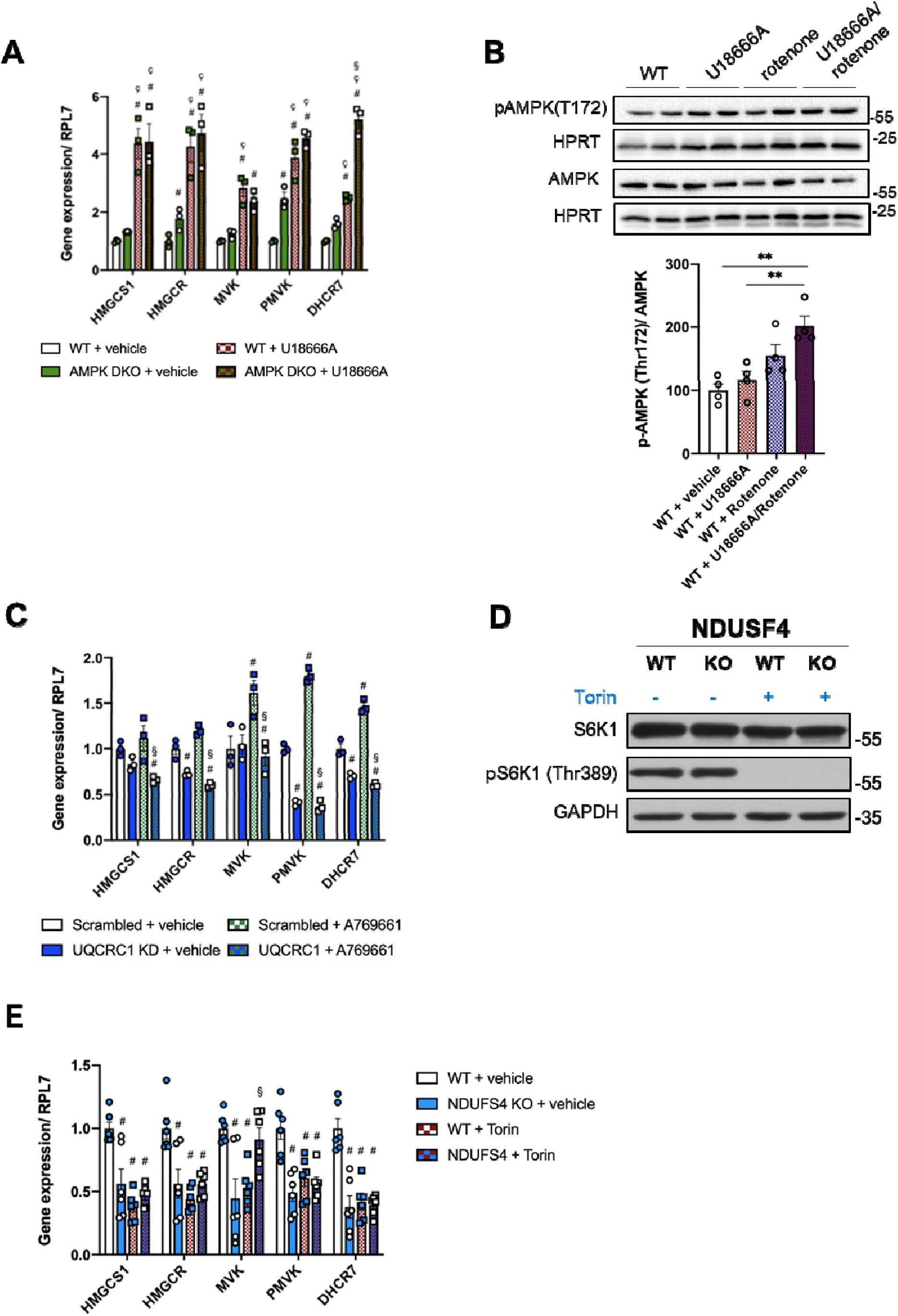
Downregulation of cholesterol synthesis in cells with mitochondrial dysfunction is independent of AMPK. **(A)** Transcript levels of cholesterol biosynthesis enzymes in WT and AMPK-DKO MEF cells, untreated or treated with U18666A. 3 independent samples were analyzed. The statistical analysis was performed using the two-way ANOVA with Tukey’s multiple comparisons test. #= significant difference from control, §= significant difference from control +U18666A. **(B)** Western blot analysis of the phosphorylated and the total form of AMPK in WT MEF cells untreated and treated with U18666A, rotenone, or U18666A + rotenone. HPRT was used as the loading control. 4 independent samples were analyzed. One way ANOVA with Tukey’s multiple comparisons test was used to determine the statistical significance. ** p < 0.001. **(C)** Transcript levels of cholesterol biosynthesis enzymes, analyzed by qPCR, in control and UQCRC1 KD HeLa cells untreated and treated with the AMPK activator A769661 (100µM, 4h). 3 independent samples were used, and the statistical significance was determined by two-way ANOVA with Tukey’s multiple comparisons test. #= significant difference from Scrambled, §= significant difference from Scrambled + A769661. **(D)** Western blot analysis of the phosphorylated (Thr389) and the total form of S6K in WT and NDUFS4 KO MEF cells, untreated and treated with the specific mTORC1 inhibitor Torin (4h, 250 nM). The same cells were analyzed through RT qPCR to evaluate the transcript levels of cholesterol biosynthesis enzymes **(E)**. The statistical significance was determined by two-way ANOVA with Tukey’s multiple comparisons test, evaluating at least 4 independent replicates #= significant difference from control, §= significant difference from NDUFS4 KO.

## SUPPLEMENTARY FILES

**Supplementary Table 1. Differentially Expressed Transcripts for the mitochondrial and lysosomal datasets.**

