## Supplementary figures and images for "Up-regulation of cholesterol synthesis by lysosomal defects requires a functional mitochondrial respiratory chain"

### Supplementary Figure 1

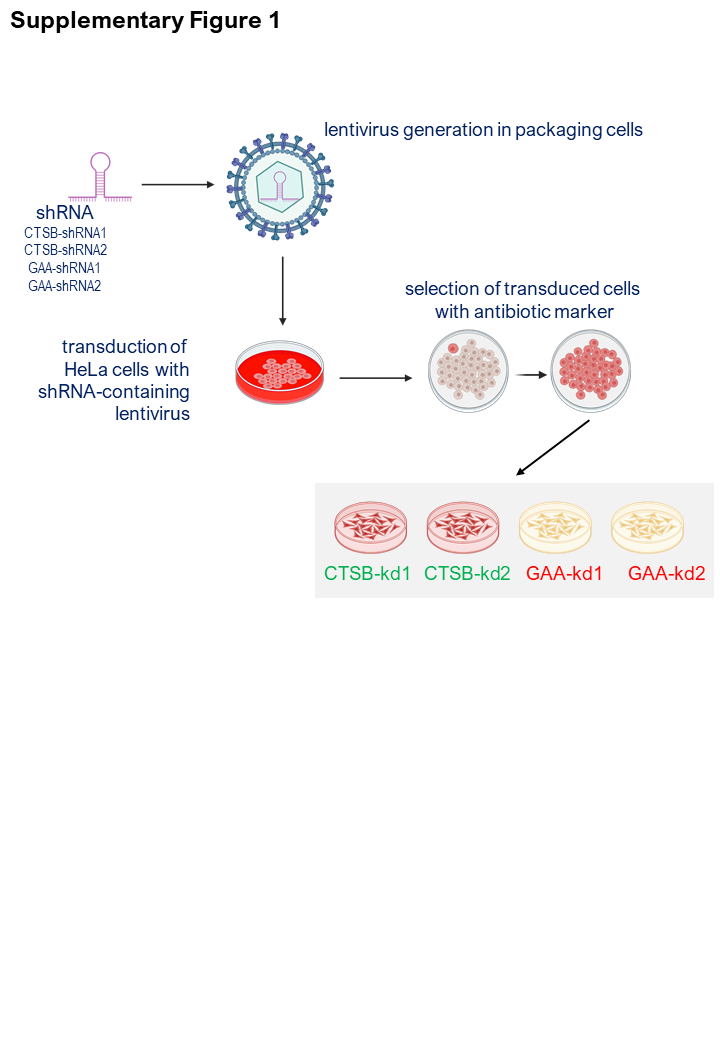

### Supplementary Figure 2

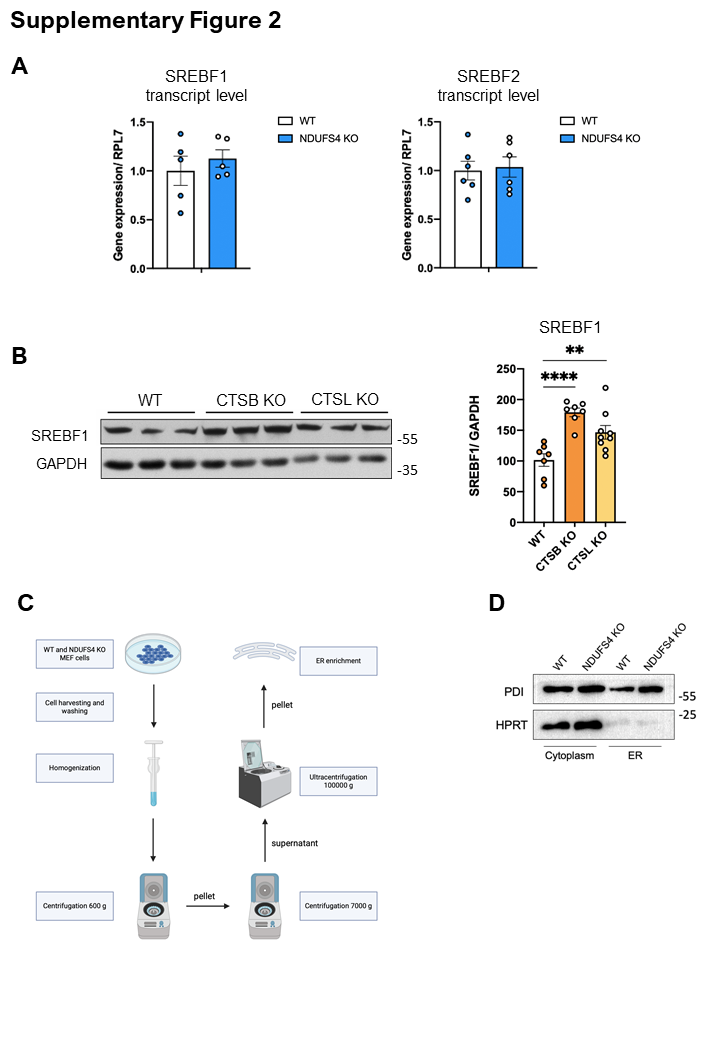

### Supplementary Figure 3

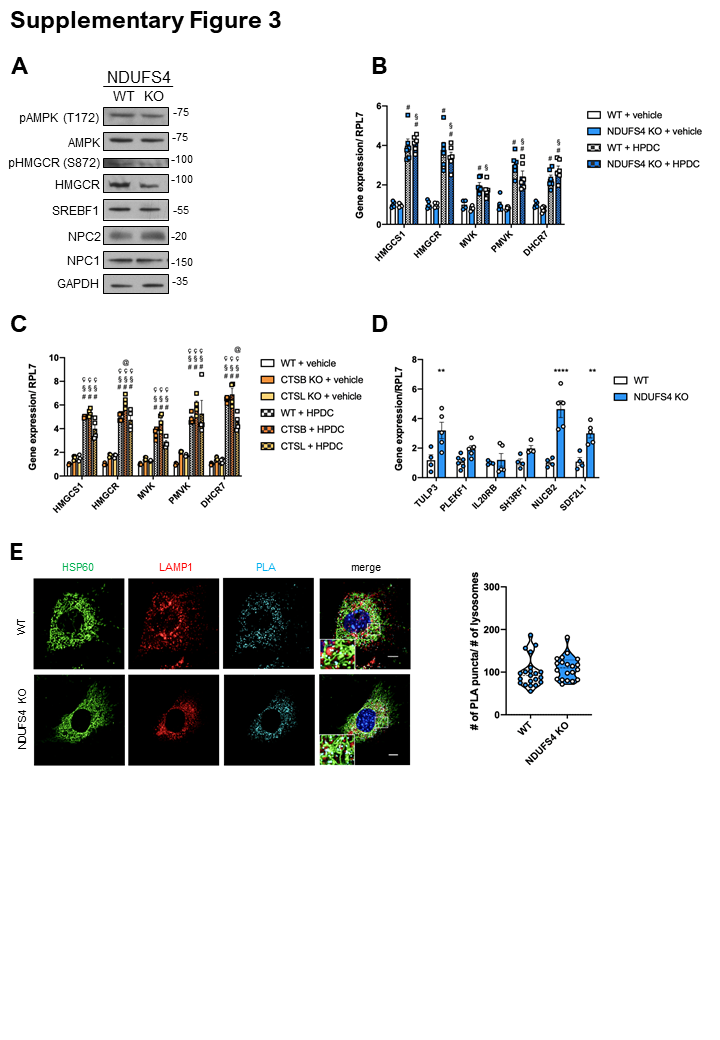

### Supplementary Figure 4

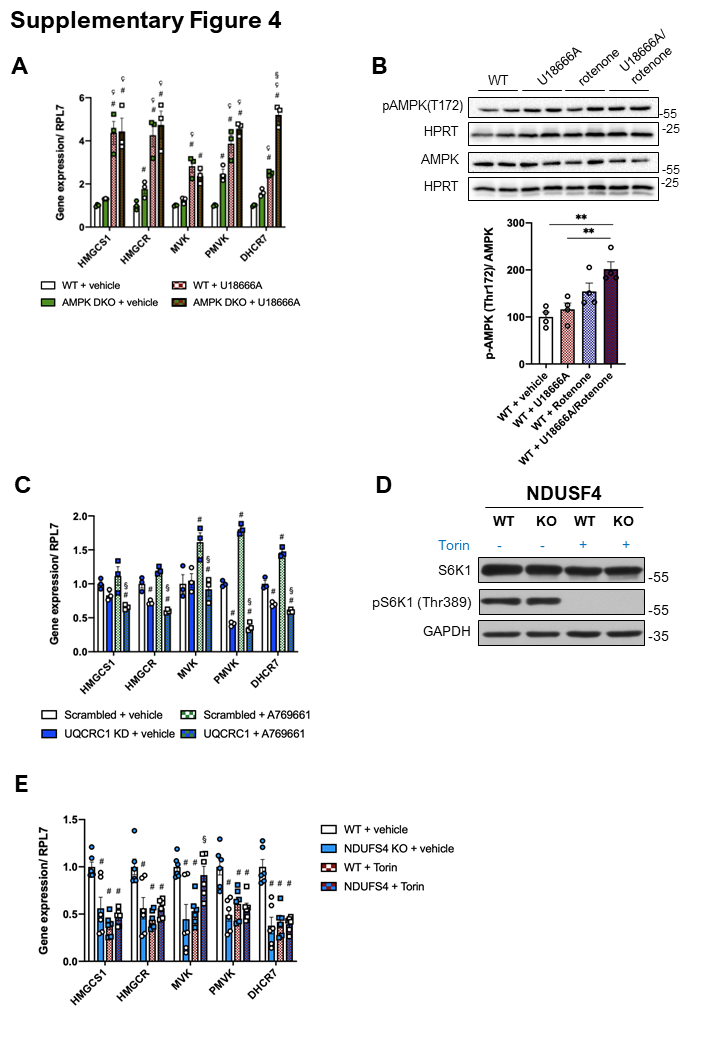
